# AhR restricts axon regeneration by balancing neuronal stress and growth response after injury

**DOI:** 10.1101/2023.11.04.565649

**Authors:** Dalia Halawani, Yiqun Wang, Jiaxi Li, Daniel Halperin, Haofei Ni, Molly Estill, Aarthi Ramakrishnan, Li Shen, Arthur Sefiani, Cédric G. Geoffroy, Roland H. Friedel, Hongyan Zou

**Author notes:** Equal Contribution (Dalia Halawani, Yiqun Wang).

## Abstract

Neurons must balance stress responses with regenerative demands following injury, but the mechanism is unclear. Here, we identify the aryl hydrocarbon receptor (AhR), a ligand-activated bHLH-PAS transcription factor, as a key regulator of this stress-growth switch. We show that ligand mediated AhR signaling restrains axon growth, whereas neuronal deletion or pharmacological inhibition of AhR promotes axonal regeneration and functional recovery in both peripheral nerve and spinal cord injury models. Mechanistic studies revealed that axotomy-induced AhR activation in dorsal root ganglion (DRG) neurons enforces proteostasis and stress-response programs to preserve tissue integrity. In contrast, AhR ablation redirects the neuronal response toward elevated de novo translation and pro-growth signaling, enabling axon regeneration. This growth-promoting effect required HIF-1α, with shared transcriptional targets enriched for metabolic and regenerative pathways. Single-cell and epigenomic analyses further revealed that the AhR regulon engages the integrated stress response (ISR) and DNA hydroxymethylation to rewire neuronal injury programs. Together, our findings establish AhR as a neuronal brake on axon regeneration, integrating environmental sensing, protein homeostasis, and metabolic signaling to control the balance between stress adaptation and axonal repair.

## INTRODUCTION

Neural injury alters the extracellular milieu due to tissue breakdown, neuroinflammation, microbial imbalance, and altered oxygen tension. To cope with stress, injured neurons deploy safeguard mechanisms that adjust metabolism or synaptic activity and activate regenerative gene programs. However, the mechanisms that balance stress adaptation with regenerative needs remain poorly understood.

Basic helix-loop-helix/PER-ARNT-SIM (bHLH-PAS) transcription factors (TFs) act as molecular sensors of environmental and physiological signals. Within the family, Bmal1 coordinates circadian rhythm, HIF-1α mediates hypoxia responses, and the aryl hydrocarbon receptor (AhR) detects xenobiotics and endogenous metabolites ^1,2^. Our recent study showed that Bmal1 gates regenerative responses of dorsal root ganglion (DRG) neurons after peripheral axotomy ^3^. Intermittent hypoxia can enhance axon regeneration through HIF activation ^4^. AhR and HIF-1α are structurally related α-subunits that share the dimerization partner ARNT (HIF-1β). The roles of AhR or ARNT in axonal injury and their interplay with HIF-1α and Bmal1 remain largely unexplored.

AhR is unique among bHLH-PAS TFs as the only ligand-activated member ^5^. Originally defined as a sensor for environmental toxins such as dioxin, AhR responds to a wide range of dietary, microbial, and metabolic molecules ^5^. Upon ligand binding, AhR translocates to the nucleus, dimerizes with ARNT, and regulates transcription of genes with AhR response elements (AHREs) ^6^. Canonical AhR signaling induces xenobiotic-metabolizing enzymes of the P450 family (CYP1A1, CYP1A2, CYP1B1) and NRF2-dependent antioxidants ^7,8^. Multiple feedback circuits restrain AhR activity, including CYP1-mediated ligand metabolism ^9^ and induction of AhR repressor (AhRR) ^10^, which competes for ARNT and represses AHRE-driven genes via chromatin remodeling ^11^. This highlights the importance of tight regulation to prevent toxicity from prolonged activation of AhR ^6^.

Competition with HIF-1α for ARNT provides another layer of control ^12,13^, though their relationship is context-dependent: AhR and HIF-1α can antagonize, cooperate, or synergize depending on temporal and cellular context ^14–17^. Recent work suggests temporally gated access to ARNT, with HIF-1α acting first and AhR subsequently taking over ^16^.

Although AhR has been extensively studied in toxicology, barrier tissue biology, and immunity, its neuronal functions, in particular in the injury setting, remain poorly defined. DRG neurons provide an excellent model to elucidate mechanisms of axon regeneration, as peripheral axotomy elicits robust axon regrowth, whereas central axotomy does not; however, a conditioning lesion (prior peripheral lesion) can prime central DRG axons for regrowth after central lesion ^18,19^.

Here, we show that DRG neurons are responsive to ligand mediated AhR signaling, which functions as a brake on axon regeneration. Neuronal deletion of AhR promoted axonal regrowth in both peripheral nerve and spinal cord injury paradigms. Mechanistic studies identified an AhR injury regulon that promotes protein homeostasis (proteostasis), while AhR loss switched neurons to elevated protein translation and metabolism, and pro-growth signaling. We further reveal a crosstalk of AhR with HIF-1α and ARNT in balancing neuronal stress adaptation and regenerative growth. These findings establish AhR as a central regulator of the stress-growth switch in response to axotomy. Our work proposes new links between environmental sensing, protein homeostasis, and axon regeneration; targeting AhR may advance therapeutic strategies for neural repair after spinal cord injury.

## RESULTS

### DRG neurons respond to ligand mediated AhR signaling

To explore transcriptional regulatory network in response to axonal injury, we analyzed a set of early-response TFs identified in axotomized DRGs at 12 or 24 h after peripheral lesion (PL, regenerative) or central lesion (CL, non-regenerative) ^20^. STRING network analysis ^21^ revealed extensive interconnections among established regeneration-associated TFs such as ATF3 ^22^, Sox11 ^23^, Jun ^24^, Smad1 ^20,25^, ATF4 ^26^, as well as immediate early genes (Egr1, Crem), but there were no functional links to AhR (**Fig. 1a**). A broader network of 39 TFs previously implicated in axon regeneration ^27^ also showed connectivity between Smad, Fos, Stat, and Jun, but no established association with AhR (**Fig. S1a**).

**Figure 1.**
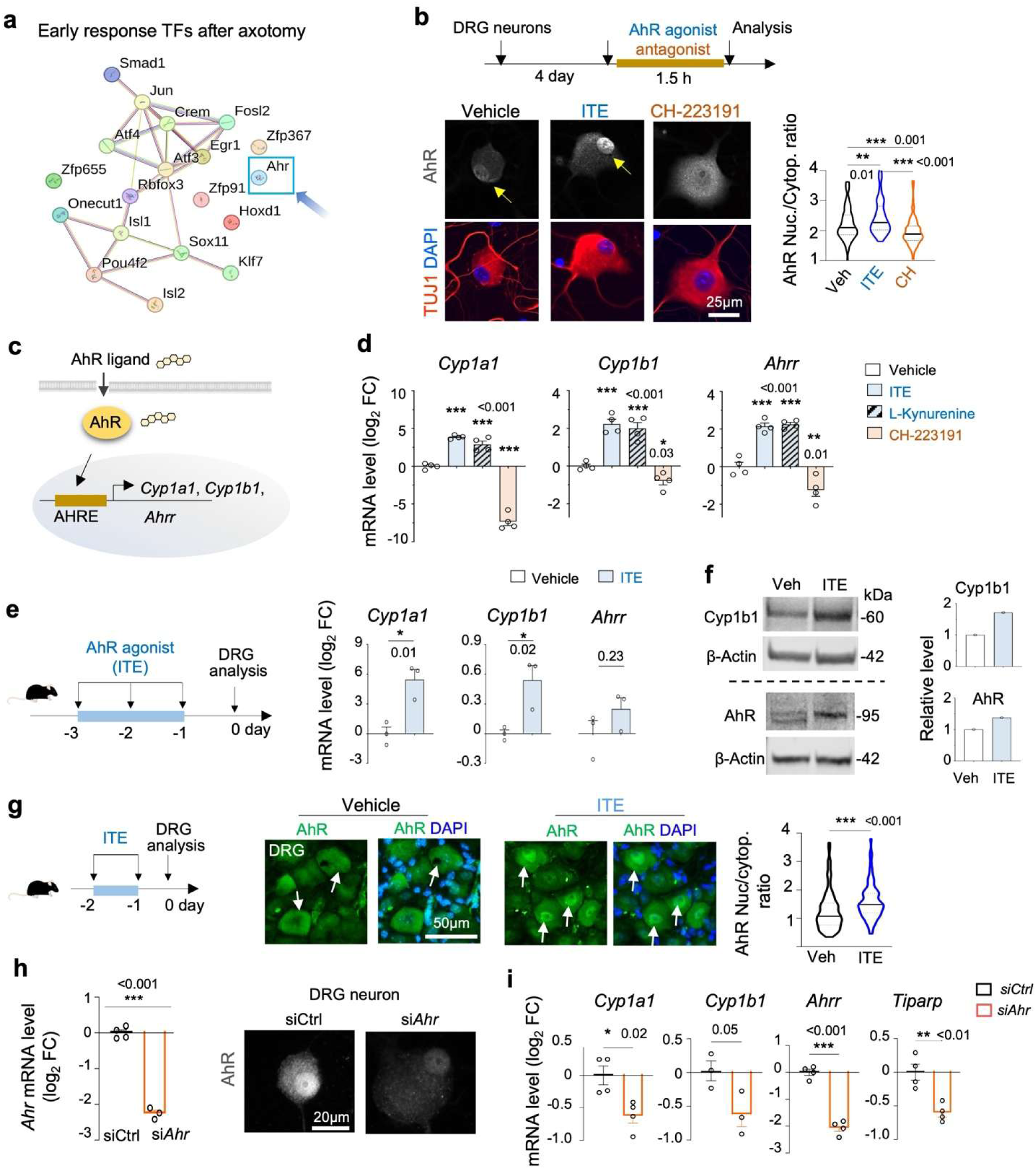
DRG neurons are responsive to ligand-mediated AhR signaling. **a.** STRING protein-protein interaction analysis revealed absence of direct signaling link between AhR and regeneration-associated transcription factors identified previously (Zou et al., 2009). **b.** Top, schematic of primary DRG neuron cultures treated with AhR agonist ITE (10 μM), antagonist CH-223191 (CH; 10 μM), or vehicle for 1.5 h before analysis. Bottom, representative IF images show cytoplasm-to-nucleus shuttling of AhR in response to ITE, while the reverse is observed with CH treatment. Quantifications of nuclear/cytoplasmic AhR ratio (n = 100 neurons per condition), with violin plots indicating median and quartiles. One-way ANOVA with Dunnett’s correction. **c.** Schematic of ligand-mediated AhR activation leading to expression of AhR response element (AHRE) containing target genes. **d.** qRT-PCR of AhR target genes in primary DRG neurons treated with vehicle, agonist (ITE; L-kynurenine) or antagonist (CH) for 24 h (25 μM drug concentrations). Values normalized to *Hprt1*. N = 4 independent cultures. Mean ± s.e.m. One-way ANOVA with Dunnett’s correction. **e.** Left, paradigm of adult mice treated with ITE (10 mg/kg i.p.) or vehicle, followed by DRG collection. Right, qRT-PCR results of AhR target genes normalized to *Hprt1* (n = 3 DRG samples from 3 mice). Mean ± s.e.m. Unpaired two-tailed Student’s t-test. **f.** Immunoblots and quantification of Cyp1b1 and AhR protein in DRGs from vehicle-or ITE-treated mice. Note electrophoretic mobility shift of AhR in ITE-treated samples. **g.** Left, paradigm of ITE treatment of adult mice, followed by DRG analysis. Right, IF analysis shows increased nuclear AhR in DRG neurons from ITE-treated mice (arrows). Quantification from 10 neurons/DRG from 9 sciatic DRGs from 3 mice per group. Violin plots show median and quartiles. Unpaired two-tailed Student’s t-test. **h.** Left, qRT-PCR analysis shows knockdown of *Ahr* by siRNA in primary DRG neurons (n = 4 cultures). Mean ± s.e.m. Unpaired two-tailed Student’s t-test. Right, IF confirms reduction of AhR protein at 48 h after onset of siRNA treatment. **i.** qRT-PCR shows downregulation of AhR target genes 48 h after knockdown of *Ahr* by siRNA (n = 4 cultures for *Cyp1a1*, *Ahrr*, and *Tiparp*; n=3 for *Cyp1b1*). Mean ± s.e.m. Unpaired two-tailed Student’s t-test.

As AhR is a ligand-activated bHLH-PAS TF, we first examined responsiveness of DRG neurons to AhR ligands. We tested stimulation by ITE, an indole-derived endogenous agonist of AhR with broad disease-modulatory roles ^28–30^, and CH-223191 (CH), a selective AhR antagonist ^31^. IF staining of cultured adult DRG neurons revealed rapid nuclear translocation of AhR upon 1.5 h stimulation by ITE, whereas CH promoted cytoplasmic retention (**Fig. 1b**). Notably, DRG neurons in primary cultures maintained in serum-free defined media displayed baseline nuclear AhR immunoreactivity even after 4 days in vitro, indicating the presence of endogenous AhR ligands (**Fig. 1b**).

As a direct readout of AhR activity, we measured gene expression of canonical AhR targets that are involved in detoxification (*Cyp1a1* and *Cyp1b1*), and *Ahrr*, a negative feedback regulator (**Fig. 1c**). qRT-PCR confirmed induction of these targets by ITE and suppression by CH (**Fig. 1d**). In vivo intraperitoneal injection of ITE for 3 days also upregulated *Cyp1a1* and *Cyp1b1* in DRGs, with a trend for increased *Ahrr* (**Fig. 1e**). Western blotting (WB) demonstrated higher Cyp1b1 protein levels in DRGs following in vivo ITE administration, and also an electrophoretic upward shift of AhR, suggesting post-translational modifications upon ligand activation (**Fig. 1f**). We also conducted IF staining for AhR in DRGs from mice treated with two daily ITE injections, which confirmed nuclear translocation of AhR into DRG neurons, with minimal signals in glial cells (**Fig. 1g**).

Next, we performed siRNA knockdown of *Ahr* in adult DRG neurons, confirmed by qRT-PCR and IF staining (**Fig. 1h**). *Ahr* silencing also resulted in downregulation of AhR targets (**Fig. 1i**). Analysis of a mouse single-cell RNA-sequencing (scRNA-seq) dataset of the nervous system ^32^ confirmed expression of *Ahr*, *Hif1a*, and *Arnt* across multiple CNS and PNS neuronal populations (**Fig. S1b**). Collectively, these findings establish that DRG neurons display responsiveness to ligand mediated AhR signaling.

### AhR inhibition promotes neurite outgrowth

To explore the function of AhR in adult DRG neurons, we conducted neurite outgrowth assays, which showed that neurons with siRNA knockdown of *Ahr* had significantly longer neurites as compared to controls (mean length of longest neurite 193 vs. 233 μm, a 21% increase; **Fig. 2a**). Concordantly, pharmacological modulation of AhR in primary adult DRG neurons revealed opposing effects, with selective antagonist CH-223191 promoting neurite elongation, whereas neurons treated with AhR agonist ITE displayed shortened neurites (**Fig. S1c**).

**Figure 2.**
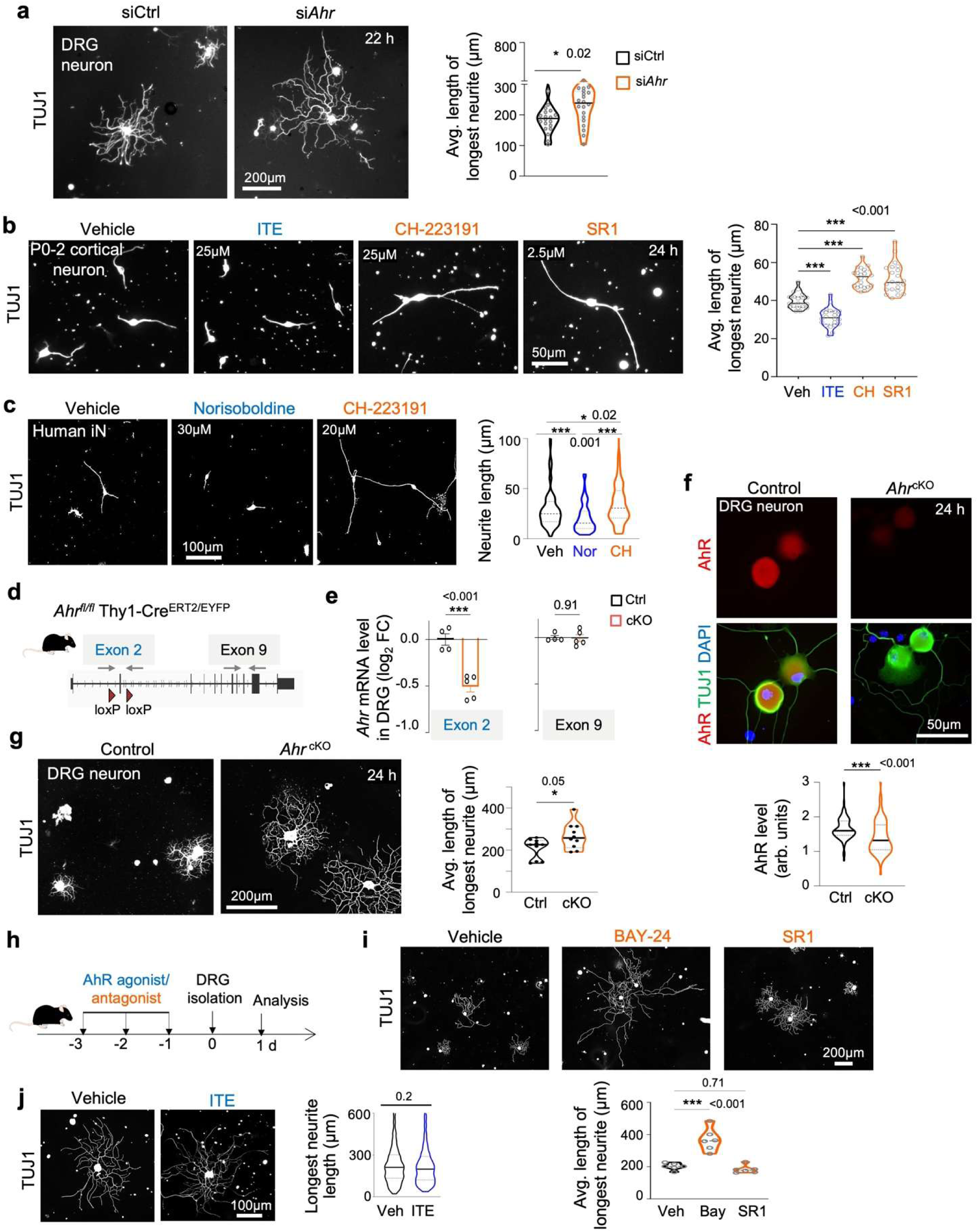
AhR inhibition enhances axon growth. **a.** Enhanced neurite outgrowth of adult DRG neurons after *Ahr* knockdown by siRNA, shown by IF for TUJ1 and quantifications of mean longest neurite length from n = 20 random fields across 4 independent cultures (each culture with more than 50 neurons each). Violin plots show median and quartiles. Mann-Whitney two-tailed test. **b.** Neonatal cortical neurons treated with AhR agonist ITE or antagonists CH-223191 or SR1. IF images and quantifications show that ITE treatment reduced neurite length, whereas CH and SR1 enhanced outgrowth. Quantification of mean longest neurite length from n = 19 random fields for Veh, 22 for ITE, 20 for CH, and 22 for SR1 across independent neuronal cultures from 2 mice. Violin plots show median and quartiles. One-way ANOVA with Dunnett’s correction. **c.** Human induced neurons treated with AhR agonist norisoboldine (Nor) or antagonist CH-223191. Nor treatment reduced neurite length, whereas CH enhanced outgrowth. Quantifications from n = 83 (Veh), 90 (Nor), and 106 (CH) cells. One-way ANOVA with Dunnett’s correction. Violin plots show median and quartiles. **d.** Diagram of *Ahr*^cKO^ allele with loxP sites flanking exon 2 and qRT-PCR primers covering exons 2 and 9. **e.** qRT-PCR shows reduced *Ahr* transcript containing exon 2 in sciatic DRGs from *Ahr*^cKO^ mice (n = 5) compared to controls (n = 4). Mean ± s.e.m. One-way ANOVA with Dunnett’s correction. **f.** IF shows reduced AhR protein levels in DRG neurons from *Ahr*^cKO^ mice compared to controls. The cKO was induced in vivo by 5 daily tamoxifen administrations (100 mg/kg, i.p.) 2 weeks before culture. Violin plots with median and quartiles, from n = 15 control and 16 cKO neurons from randomly selected fields of 2 independent cultures. Mann-Whitney two-tailed test **g.** IF shows enhanced neurite outgrowth of DRG neurons from *Ahr*^cKO^ mice compared to littermate controls. Quantifications of mean longest neurite length from n = 10 random fields across 2 independent cultures from n = 4 mice per genotype. Violin plots with median and quartiles. Mann-Whitney two-tailed test. **h-j.** Experimental paradigm and IF images of DRG neurons from mice treated with AhR agonist ITE, antagonists BAY-24 or SR1, or vehicle. For BAY-24 and SR1 study, quantification of mean longest neurite length from n = 6 randomly selected fields across independent cultures. One-way ANOVA with Dunnett’s correction. Violin plots show neurite length distribution with median and quartiles. Neurite outgrowth was measured one day after plating. For ITE study, n = 466 (vehicle) and 533 (ITE) neurons from n = 3 mice for each condition. Unpaired two-tailed Student’s t-test.

We also tested AhR effect in CNS neurons. In mouse neonatal cortical neurons and human induced neurons (iNs) derived from embryonic stem cells, we observed similar growth-promoting or inhibitory effects of AhR antagonists and agonists, respectively (**Fig. 2b, c**). A broader screen of AhR antagonists/agonists with human iNs confirmed these effects in a dose-dependent manner (**Fig. S1d**). Further supporting these findings, the AhR antagonist StemRegenin1 (SR1) was independently identified in an unbiased high-throughput screen for compounds that enhance neurite outgrowth of mouse adult cortical neurons ^33^. SR1 increased neurite length by 28% and neurite initiation by 58%, compared to vehicle (**Fig. S1e**).

To further corroborate the role of AhR in neurons, we generated tamoxifen-inducible neuronal conditional *Ahr* knockout (cKO) mice by crossing *Ahr^fl/fl^* and Thy1-CreERT2/EYFP (SLICK-H) strains ^34,35^ (**Fig. 2d**). We first confirmed wide expression of EYFP in DRG neurons and sciatic axons (**Fig. S2a**). Breeding of SLICK-H mice with the Rosa26-LSL-Sun1-GFP (INTACT) reporter line ^36^ confirmed efficient neuronal tagging in DRG neurons upon tamoxifen injection (**Fig. S2b**).

Genotyping demonstrated excision of exon 2 of *Ahr* floxed allele in DRG, but not in tail tissues, two weeks after tamoxifen administration (**Fig. S2c, d**). qRT-PCR of DRGs showed reduction of *Ahr* exon 2 but not exon 9 containing transcripts (**Fig. 2e**). RNA-seq data analysis similarly confirmed the selective loss of *Ahr* exon 2 reads in DRG of *Ahr*^cKO^ mice, while distal exons remained intact (**Fig. S2e**). IF staining validated loss of nuclear AhR immunosignals in cultured DRG neurons from *Ahr*^cKO^ mice (**Fig. 2f**). Likewise, AhR protein was largely absent in DRG neurons of ITE-treated *Ahr*^cKO^ mice (**Fig. S2f**). Notably, AhR protein levels in DRG neurons were higher than in glial cells (**Fig. S2g**).

Functionally, *Ahr*^cKO^ DRG neurons extended significantly longer neurites than controls at 24 h post-seeding (mean length of longest neurite 393 vs. 232 μm, a 70% increase) (**Fig. 2g**), corroborating the results from *Ahr* knockdown and pharmacological inhibition.

We then explored the in vivo efficacy of AhR antagonists SR1 and Bay2416942 (Bay), both in clinical development ^37^. Interestingly, in vivo intraperitoneal injection of Bay but not SR1 for 3 days resulted in longer neurites of seeded DRG neurons, reflective of specific in vivo pharmacokinetics/dynamics of different compounds (**Fig. 2h, i**). The in vivo treatment with AhR agonist ITE did not cause shorter neurites in cultured DRG neurons (**Fig. 2j**), consistent with the self-limiting nature of AhR signaling by multiple negative feedback loops ^6^.

Together, both genetic and pharmacological approaches demonstrate that AhR activity restrains neurite growth, whereas AhR inhibition or genetic ablation can promote axon elongation across species and neuronal subtypes.

### PL triggers an early induction of AhR in axotomized DRGs

To understand AhR signaling in the conditioning lesion paradigm, we conducted RNA-seq analysis on axotomized DRGs after PL, using the contralateral DRGs from the same mice to control for anesthesia, pain or other systemic effects (**Fig. 3a**). We identified a positive enrichment for both xenobiotic metabolism and hypoxia pathways after PL (**Fig. 3b**). Consistently, mRNA reads for *Ahr, Hif1a*, *Arnt* and AhR targets *Cyp1b1* and *Tiparp* ^38^ were all elevated after PL (**Fig. 3c**; **Table S1**). *Cyp1a1* and *Ahrr* showed lower expression levels in sciatic DRG.

**Figure 3.**
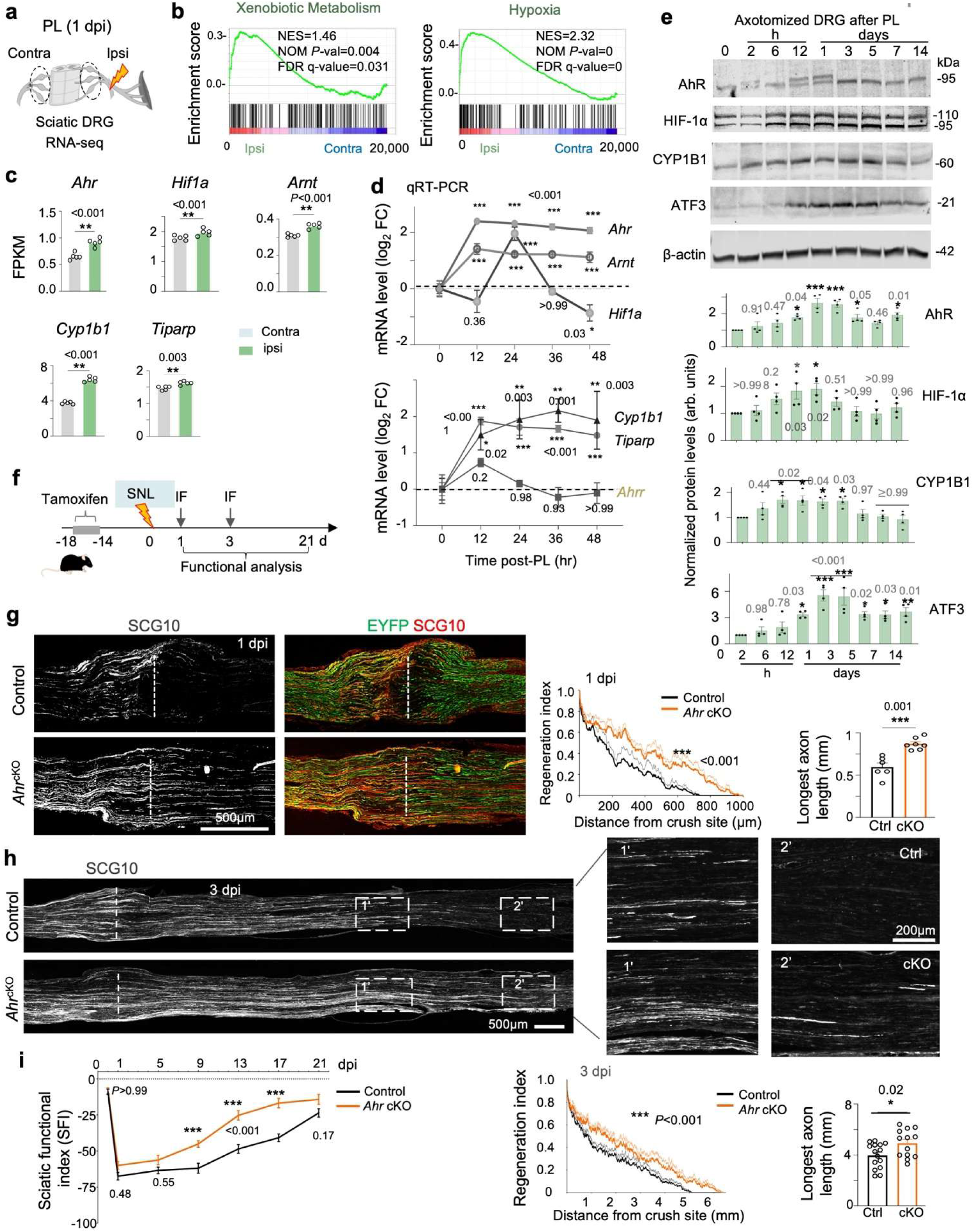
Neuronal AhR deletion enhances axon regeneration after sciatic nerve injury. **a.** Experimental design of RNA-seq to compare ipsilateral and contralateral sciatic DRGs (L4-6) of wild-type mice at 1 dpi after peripheral nerve lesion (PL). **b.** GSEA shows enrichment of xenobiotic metabolism and hypoxia pathways in ips-vs. contralateral DRGs after PL. NES, normalized enrichment score; FDR, false discovery rate. **c.** mRNA reads in FPKM (fragments per kb transcript per million reads) for the indicated genes in ipsi- and contralateral DRGs at 1 dpi. n = 5 DRG samples per condition. Mean ± s.e.m. Unpaired two-tailed Student’s t-test. **d.** Time-course analysis of transcription of *Ahr, Arnt, Hif1a* and of AhR downstream target by qRT-PCR in DRGs after PL, normalized to *Hprt1*. n = 3 mice, each with pooled L4-6 DRGs per timepoint. Mean ± s.e.m. One-way ANOVA with Dunnett’s correction. **e.** Immunoblots and quantification of protein levels for AhR, HIF-1α, Cyp1b, and ATF3 in DRGs at successive time points after PL. β-actin as loading control. n = 4 mice per time point. Mean ± s.e.m. One-way ANOVA with Dunnett’s correction. **f.** Experimental scheme of sciatic nerve lesion (SNL) and subsequent analysis. **g, h.** IF images and quantifications of SCG10^+^ regenerating axons traversing the lesion site (dashed line indicates lesion center) at 1 dpi and 3 dpi. n = 6 control and 7 cKO mice at 1 dpi; n = 15 control and 13 cKO mice at 3 dpi. Regeneration index: two-way ANOVA with Bonferroni correction. Maximal axon length distal to lesion center: unpaired two-tailed Student’s t-test. Mean ± s.e.m. **i.** Sciatic functional index (SFI) in control and *Ahr* cKO mice across time, showing enhanced recovery in cKO at 9, 13, and 17 dpi. Two-way ANOVA with Bonferroni multiple test correction. Mean ± s.e.m.

Time-course qRT-PCR confirmed early induction of *Ahr* and *Arnt* at 12 h post-PL in axotomized DRGs, persisting to 48 h. By contrast, *Hif1a* was induced with a slight delay, peaking at 24 h and returning to baseline by 48 h (**Fig. 3d**). *Cyp1b1* and *Tiparp* were similarly upregulated, while *Ahrr* remained unchanged. Analysis of a single-cell RNA-seq dataset ^39^ further verified induction of *Ahr, Hif1a*, and *Arnt* across major DRG neuron subtypes from 12 to 72 h post-PL (**Fig. S2h**).

WB corroborated these findings, showing AhR protein upregulation in DRGs as early as 2 h post-PL, increasing by 12 h, peaking at 1-3 dpi, and persisting at 14 dpi, paralleling ATF3 expression, a marker of conditioning lesion ^22^ (**Fig. 3e**). WB for AhR also exhibited an upward band shift after PL, particularly at 12-24 h (**Fig. 3e**), similar to that induced by AhR agonist. HIF-1α protein was induced transiently, detectable at 12 h, peaking at 1 dpi, and declining by 3 dpi (**Fig. 3e**). In line with heightened AhR activity, CYP1B1 showed transient protein induction at 1-3 dpi. Thus, PL evokes an early but transient AhR activation despite persistent AhR expression, consistent with feedback mechanisms that rapidly attenuate signaling ^6^.

### AhR inhibition enhances in vivo axonal regeneration after sciatic nerve and spinal cord injury

We next examined the in vivo effects of neuronal *Ahr* deletion on axon regeneration in the sciatic nerve lesion (SNL) paradigm. IF staining of nerve sections for SCG10, a marker of regenerating sensory axons ^40^, revealed in *Ahr*^cKO^ mice significantly more and longer axons extending beyond the crush site compared to controls at both 1 dpi (869 vs. 594 μm, ∼50% increase) and 3 dpi (5.8 vs. 4.8 mm, ∼20% increase) (**Fig. 3f-h**). The regenerative phenotypes seen at 1 and 3 dpi signify that *Ahr* deletion primes DRG neurons for a rapid activation of pro-repair program upon injury.

Following sciatic nerve crush injury, both genotypes initially exhibited toe flexion deficits and weight-bearing failure, but from 9-21 dpi, *Ahr*^cKO^ mice showed significantly improved sciatic functional index (SFI) (**Fig. 3i**). Of note, neuronal AhR loss did not impair baseline motosensory performance: ladder walking (regular and irregular rung spacing) and tactile sensitivity (von Frey filament assay) were similar between *Ahr*^cKO^ and control mice after both short-(2-5 weeks) and long-term (14 months) AhR deletion (**Fig. S2i**).

We confirmed the peripheral nerve regeneration results in *Ahr*^fl/f*l*^ *Nestin*^Cre^ mice ^41,42^ as an independent model, with near ablation of AhR proteins in DRG and brain tissues by WB (**Fig. S3a**). WB also demonstrated robust AhR induction in axotomized DRG at 3 dpi, which was blunted in Nestin-Cre cKO though still mildly elevated compared to naive state, suggesting glial AhR expression (**Fig. S3b**). After sciatic crush, Nestin-Cre cKO mice exhibited more and longer SCG10^+^ axons beyond the lesion at 2 dpi, reflected in higher regenerative index and maximal axonal length (1.44 vs. 1.06 mm, +36%) (**Fig. S3c**). Functional assays confirmed accelerated recovery, with earlier toe spreading and higher SFI from 9-21 dpi (**Fig. S3d**), accompanied by enhanced hindpaw reinnervation (PGP9.5^+^ sensory axons at 21 dpi) (**Fig. S3e**). Importantly, Nestin-Cre *Ahr*^cKO^ mice remained viable and fertile with no baseline sensorimotor deficits (**Fig. S3f**), unlike constitutive <u>AhR</u> knockouts that develop demyelination and inflammation phenotypes ^43,44^.

We next examined the SNL recovery of older *Ahr*^cKO^ mice (cKO induction with Thy1-CreER/EGFP 14 months before injury), which confirmed that a pro-regenerative effect of *Ahr* deletion on peripheral nerve regeneration was maintained in aged animals (**Fig. S4a, b**). However, when AhR deletion or pharmacological inhibition with SR1 or Bay was applied after SNL, no enhanced axon regrowth was observed (**Fig. S4c, d**), indicating the importance of an early priming effect of AhR ablation to achieve additional gain over the already robust regenerative capacity of DRG neurons after peripheral nerve injury.

We next asked whether neuronal AhR ablation could promote axon regrowth and functional recovery in the CNS injury setting where regeneration is minimal (**Fig. 4a**). We applied thoracic contusion spinal cord injury (SCI) with an impactor device at T8 level, which yielded comparable force and tissue displacement parameters of injury severity in control and *Ahr*^cKO^ mice (**Fig. 4b**). IF staining of spinal cord sections at 35 dpi for neurofilament H (NFH) revealed substantially more axon bundles traversing the dorsal spinal cord in *Ahr*^cKO^ mice relative to controls (**Fig. 4c**). Consistent with these histological findings, cKO mice also demonstrated superior motor-sensory recovery through 5-week post-injury period, reflected by higher Basso Mouse Scale (BMS) scores in open-field locomotion, fewer errors on ladder walking across both rung patterns, and improved tactile sensitivity in von Frey testing (**Fig. 4d, e**; **Video S1-S6**). Neuronal *Ahr* deletion did not alter lesion size demarcated by GFAP and CSPG staining (**Fig. 4f**).

**Figure 4.**
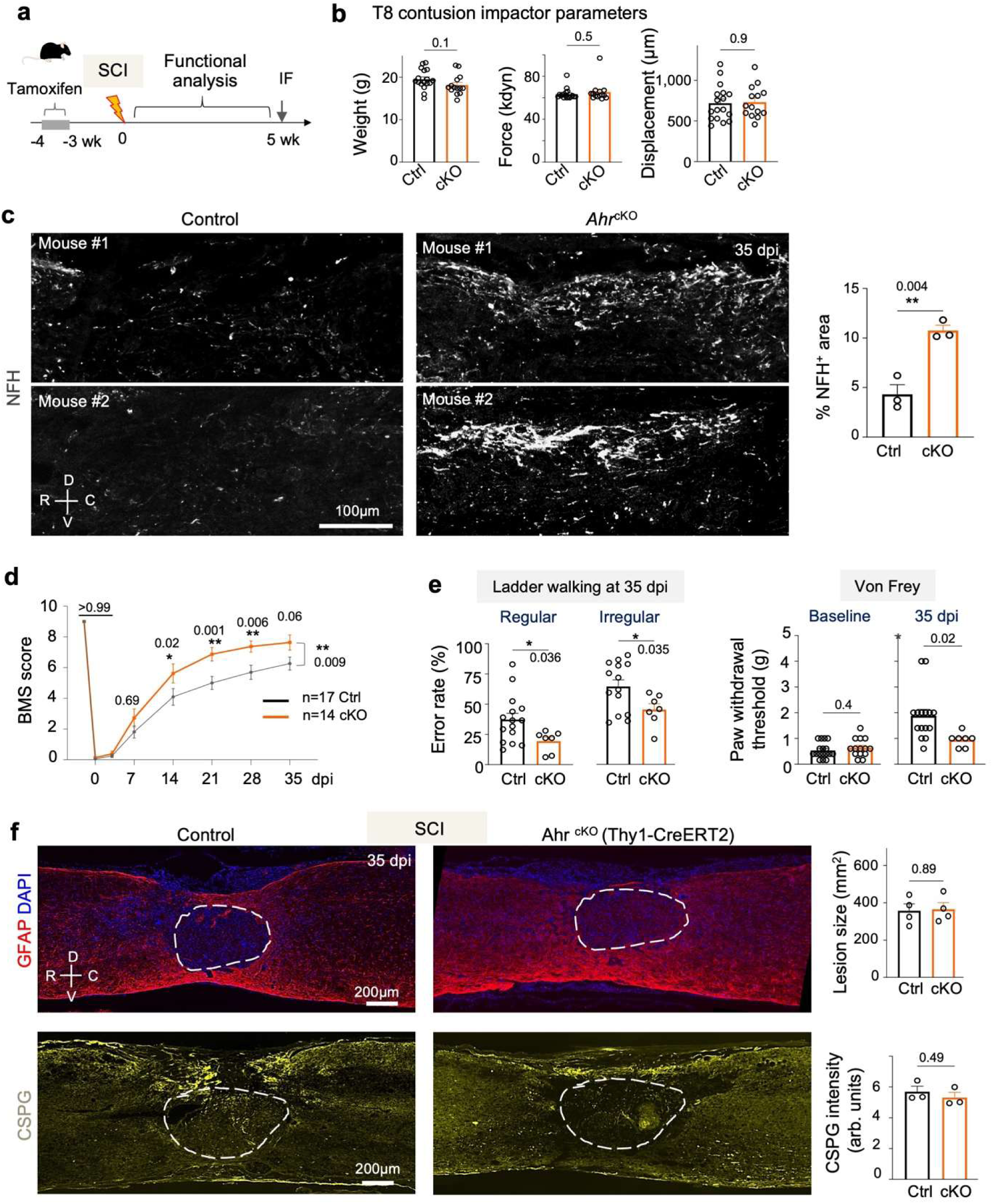
Neuronal AhR deletion enhances axon regrowth and functional recovery after SCI. **a.** Experimental design. Tamoxifen (100 mg/kg, i.p., daily for 5 days) was administered 4 weeks before T8 contusion injury for both *Ahr*^cKO^ (with Thy1-CreER/EYFP) and control cohorts. **b.** Measurements of contusion injury parameters with impactor device in cKO and control cohorts. For impactor parameters: weight, Force, and displacement, n = 17 control and 14 cKO mice. Mean ± s.e.m. Unpaired two-tailed Student’s *t*-test. **c.** IF for neurofilament-H (NFH) shows an increase of axons traversing lesion site in cKO mice compared to controls at 35 dpi. n = 3 mice per group. Mean ± s.e.m. Unpaired two-tailed Student’s t-test. **d.** Locomotor performance scoring with Basso Mouse Scale (BMS) demonstrates improved recovery in cKO mice. Average BMS score of n = 17 control and 14 cKO mice per datapoint. Mean ± s.e.m. Two-way ANOVA with Tukey’s correction. **e.** Ladder walking error rates (with regular and irregular spacing of rungs) at 35 dpi and von Frey filament response thresholds before injury (baseline) and at 35 dpi. cKO mice showed improved motor coordination and sensory recovery. n = 15 for control and 7 for cKO mice for regular ladder and von Frey assays; n=14 for control and 7 for cKO mice for irregular rung ladder. Mean ± s.e.m. Unpaired two-tailed Student’s t-test. **f.** IF for reactive astrocyte marker GFAP and matrix proteoglycan CSPG (stained with antibody clone CS-56) at 35 dpi after SCI show comparable size of lesion between cKO and control mice. n = 4 mice per group for GFAP and n=3 mice per group for CSPG analysis. Mean ± s.e.m. Unpaired two-tailed Student’s t-test.

As a proof-of-principle for translational application for SCI therapy, we also treated a cohort of mice with AhR inhibitor SR1 after the onset of SCI (**Fig. S5a, b**). Encouragingly, when compared to vehicle controls, SR1 treated animals showed significantly improved motor-sensory functional recovery (BMS scoring) as early as 14 dpi and sustained through 21 to 35 dpi (**Fig. S5c**). This was also paralleled by improved performance of SR1 treated animals in ladder walking and von Frey filament assays at 35 dpi (**Fig. S5d**; **Video S7-S12**). Together, these results demonstrate that neuronal AhR ablation enhances axonal regrowth and functional recovery after both peripheral nerve lesion and SCI.

### Gut microbial deletion does not affect axon regeneration, and neuronal AhR ablation does not alter immune responses after injury

As AhR stability and cytoplasmic retention are regulated by HSP90, XAP2 (*Aip*), and p23 (*Ptges3*) (**Fig. S6a**), we also examined their expression after PL. RNA-seq analysis of DRG transcriptomes revealed modest changes of these genes at 1 dpi, while qRT-PCR detected reduction of *Aip* and *Ptges3* at 36 h (**Fig. S6b, c**), supporting early activation of AhR after PL.

We also examined involvement of L-kynurenine pathway in conditioning lesion, as a ligand source for AhR activation ^45^ (**Fig. S6d**). Analysis of RNA-seq data revealed limited changes in key biosynthetic enzymes that convert tryptophan to L-Kyn or kynurenic acid at 1 dpi after PL, with *Tdo2* showing slight upregulation, while *Kyat3* showed a modest down-regulation (**Fig. S6e**). Of note, mRNA reads for *Ido1/2* and *Tdo2* were low in DRGs. qRT-PCR analysis detected downregulation of *Kyat1* and no significant change for *Kmo* in PL DRGs at 36 h (**Fig. S6f**). Collectively, these results indicate no major transcriptional changes of the enzymes involved in L-kynurenine pathway in axotomized DRGs.

Given that microbial metabolites can act as AhR ligands ^46^, we depleted the gut microbiome with antibiotics ^47^ before SNL (**Fig. S7a**). We observed no difference in DRG neurite outgrowth in culture or in in vivo after nerve injury between control and antibiotics treated groups (**Fig. S7b, c**). Indole-3-propionate (IPA) is a gut microbiota-derived metabolite that is linked to axon regeneration through neutrophil chemotaxis ^48^; it can also act as an AhR ligand to modulate sepsis and neuroinflammation ^49,50^, although the serum concentration of IPA is far below the levels needed to produce an AhR-dependent physiological effect ^51^. We tested IPA treatment of DRG neurons in culture, which resulted in no improvement on DRG neurite outgrowth, and at higher concentrations even reduction in outgrowth (**Fig. S7d**).

As the Thy1-CreERT2/EYFP allele of our cKO mice is also expressed in enteric neurons, we examined immune composition in the gut epithelium, but detected no significant changes (**Fig. S8a-c**). Likewise, neuronal AhR ablation had no overt effects on immune responses to peripheral nerve injury. In sciatic DRGs, IF showed no differences in CD45^+^ leukocytes, CD206^+^ anti-inflammatory myeloid cells, CD68^+^ phagocytes, IBA1^+^ macrophages, or PU.1^+^ myeloid lineage cells under naive or axotomized conditions (**Fig. S9a, b**). Other immune subsets, including NK1.1^+^ NK cells, Ly6G^+^ neutrophils, and CD4^+^ or CD8^+^ T cells, were rarely detected in DRGs and were unaffected by neuronal *Ahr* deletion. At the sciatic nerve crush site, both the density and spatial distribution of F4/80^+^ macrophages and Sox10^+^ Schwann cells were comparable between *Ahr*^cKO^ and control mice (**Fig. S9c**). These results suggest that the enhanced axonal regrowth detected in *Ahr*^cKO^ mice arises from a neuron-intrinsic mechanism rather than altered immune or Schwann cell responses.

### AhR-dependent transcriptomic changes after PL highlight translational regulation and proteostasis

To assess the pathways impacted by neuronal deletion of AhR in axonal injury condition, we profiled DRG transcriptomes from *Ahr*^cKO^ and control mice at 1 dpi after PL (n = 5 per condition), using contralateral DRGs for comparison to control for systemic influences such as anesthesia, surgical stress, or pain (**Fig. 5a**; **Table S1, S2**). Principal component analysis revealed that injury status, rather than genotype, accounted for the largest variance (PC1, 43%), with ipsi- and contralateral DRGs forming distinct clusters, whereas genotype did not segregate samples either before or after injury (**Fig. S10a**). Thus, *Ahr* cKO did not broadly reprogram DRG transcriptome but induced pathway-specific shifts.

**Figure 5.**
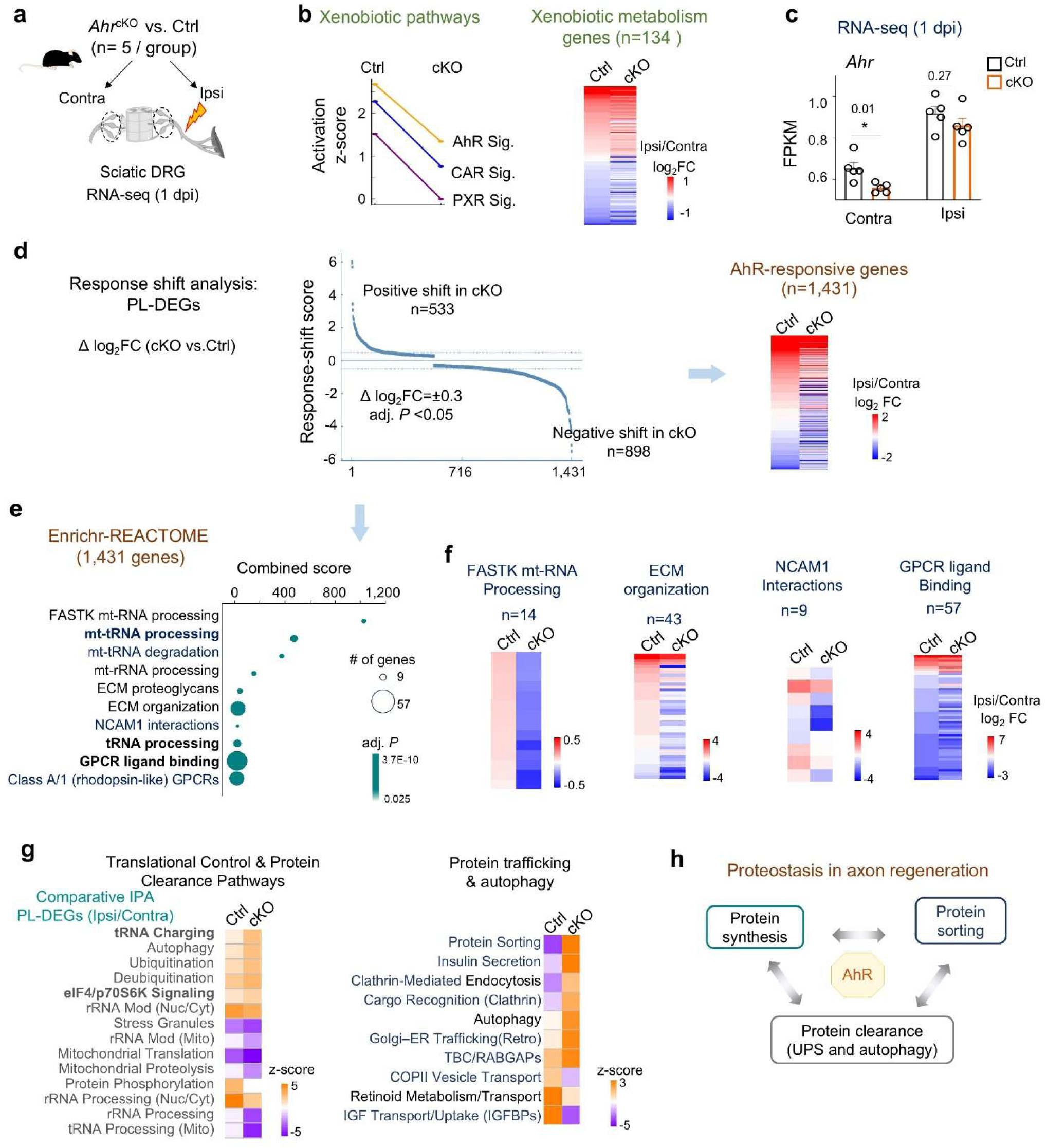
AhR controls proteostasis pathways in response to axotomy. **a.** Experimental design of RNA-seq of ipsi- and contralateral L4-L6 DRGs from *Ahr*^cKO^ and littermate controls at 1 dpi (n = 5 per group). **b.** Xenobiotic metabolism pathways were suppressed in cKO compared to controls. Heatmap shows AhR-dependent response shifts in 134 xenobiotic metabolism genes after PL. **c.** RNA-seq analysis revealed reduced *Ahr* expression (fragments per kb transcript per million reads; FPKM) in cKO versus control DRGs. Note robust *Ahr* induction in ipsilateral DRGs after PL. **d.** Response shift score (log_2_FC[cKO]) - log_2_FC[Ctrl]) across PL-DEGs. Heatmap shows changes for 1,431 AhR-responsive genes at 1 dpi after PL. **e.** Enrichr pathway analysis of AhR-responsive genes after PL. **f.** Heatmaps show gene expression changes in the enriched pathways between control and cKO after PL. **g.** Comparative IPA of PL-DEGs (ipsi/contra) highlighting enriched pathways related to translational control and proteostasis. **h.** Model of AhR-dependent proteostasis regulation in axon regeneration.

Comparison of ipsi-vs. contralateral DRGs identified 2,658 PL-DEGs in cKO and 2,501 in control, with 1,802 overlapping genes showing largely concordant directionality, though some displayed amplitude shifts or divergent responses (**Fig. S10b, c**; **Table S3**). Hence, neuronal *Ahr* deletion resulted in a selective rather than global alterations of the PL-associated transcriptome. Canonical xenobiotic programs (AhR, PXR, CAR) were selectively suppressed in *Ahr*^cKO^ DRGs, with attenuated induction of 134 curated xenobiotic genes (**Fig. 5b**; **Table S4**). Regarding *Ahr* mRNA expression, in controls, *Ahr* was robustly upregulated in ipsilateral DRG at 1 dpi; in cKO, *Ahr* were reduced, more prominently in contralateral (log_2_FC = −0.24, adj *P* = 0.02) than in ipsilateral (log_2_FC = −0.10, adj *P* = 0.13), reflecting glial induction after PL (**Fig. 5c**).

To capture broader AhR-dependent effects, we calculated response-shift scores (RSS; log_2_FC[cKO] – log_2_FC[Ctrl]) across PL-DEGs (adj *P* ≤ 0.05 in either genotype). Applying a threshold of |RSS| ≥ 0.3, we identified 1,431 AhR-responsive genes (898 negative and 533 positive RSS; **Fig. 5d**; **Table S5**). Enrichment analysis highlighted pathways associated with translational control, including rRNA, mitochondrial tRNA processing, and ECM interactions, with most genes showing negative RSS, consistent with the role of AhR as a transcriptional activator (**Fig. 5e, f**). GPCR signaling was also enriched, which can modify translational machinery, leading to rapid changes in protein synthesis within minutes ^52^.

We next analyzed the AhR-responsive genes by applying ChEA3 analysis, an integrative TF enrichment analysis tool that combines ChIP-seq datasets, gene co-expression, and TF-gene co-occurrence ^53^. This revealed 132 putative direct AhR targets, of which 87% displayed negative RSS in cKO compared to control (**Fig. S10d**; **Table S6**). These genes were strongly enriched in translational pathways, anchoring the broader AhR-dependent signatures. Additional enriched pathways included BDNF, HIF, senescence, autophagy, and various metabolism such as Vitamin B12, selenium, folate, LDL (**Fig. S10e**).

Comparative IPA of PL-DEGs (ipsi/contra) between genotypes revealed that control DRGs preferentially engaged proteostasis pathways, including ribosomal RNA processing and modification (linked to ribosome assembly), mitochondrial translation, break down or clearance of metabolites, proteins, and organelles (leucine degradation, pexophagy, mitochondrial proteolysis, and glutaryl-CoA degradation, mRNA decay) (**Fig. S10f-h**). In contrast, *Ahr*^cKO^ DRGs activated growth-promoting signaling (Neurexin, NGF, insulin, PDGF, L1CAM), protein sorting, and histone modification together with lipid biosynthesis. Closer inspection of proteostasis pathways suggested mitochondrial-cytosolic crosstalk, with cytosolic tRNA charging and ubiquitination upregulated while mitochondrial pathways suppressed in cKO (**Fig. 5g**). Protein trafficking pathways, including autophagy and endocytosis, were also activated (**Fig. 5g**). Thus, AhR upregulation after axotomy may function to reinforce protein quality control and proteostasis, whereas its loss reprograms the injury response toward biosynthesis and pro-growth signaling (**Fig. 5h**).

### AhR cKO enhances de novo protein synthesis in axotomized DRG neurons

To assess directly the impact of AhR ablation on neuronal protein synthesis, we conducted puromycin (Puro; a tyrosyl-tRNA analog) incorporation assay, which labels elongating peptides (**Fig. 6a**). We validated the assay by demonstrating robust, dose-dependent signals in cultured DRG neurons following puromycin labeling (**Fig. 6b**). Notably, we observed stronger Puro incorporation in neurons than in glia, signifying higher neuronal translational activity (**Fig. 6b**).

**Figure 6.**
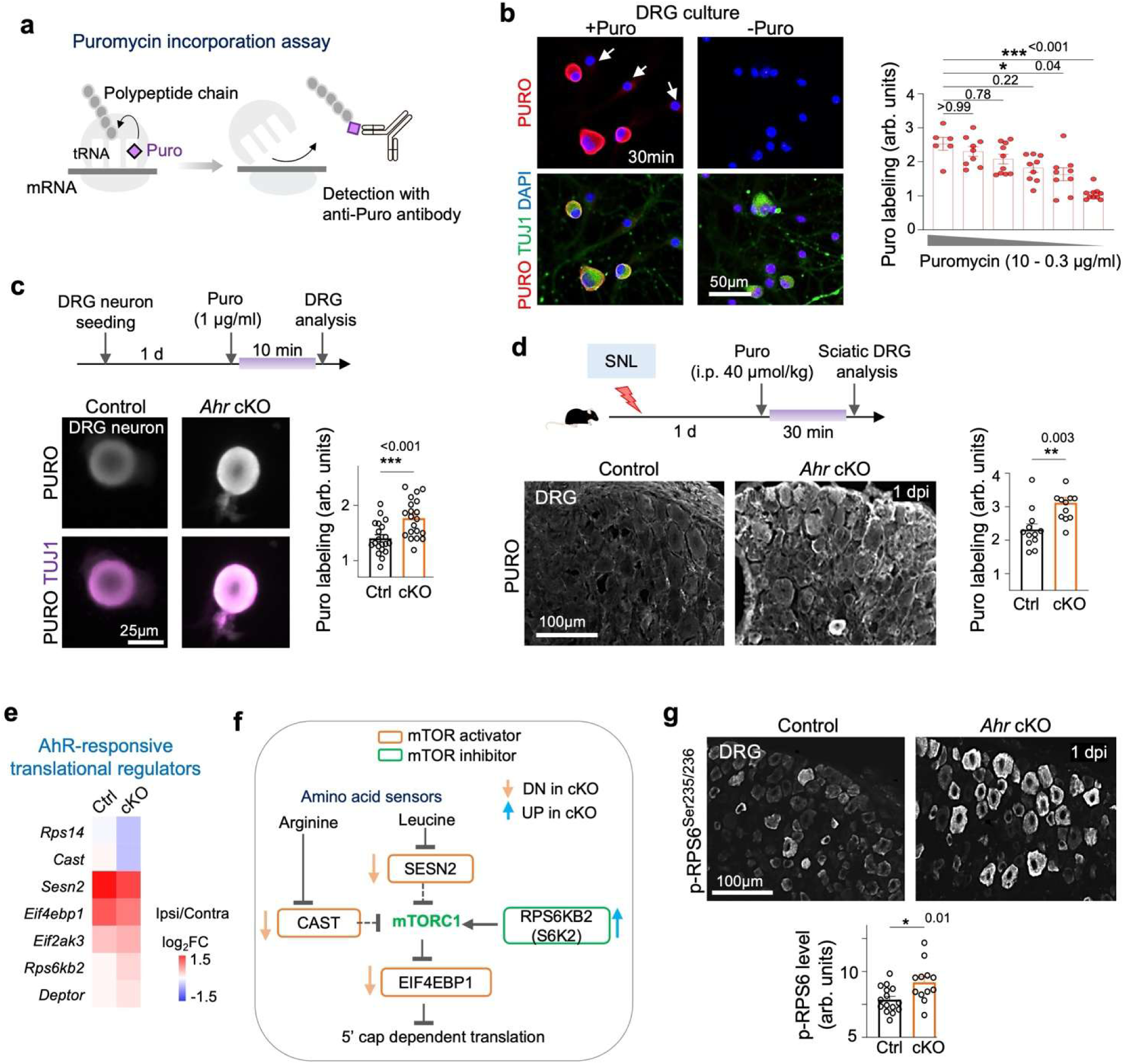
Neuronal AhR ablation enhances global protein synthesis in response to axotomy. **a.** Schematic of puromycin incorporation assay to assess de novo protein synthesis. **b.** Validation of assay in DRG cultures. IF images show absence of labeling without puromycin, but robust labeling in TUJ1^+^ neurons after 30 min puromycin exposure, with low labeling in TUJ1^-^ glia (arrows). Quantifications demonstrate dose-dependent incorporation. Average puromycin labeling intensity from n = 10 randomly selected fields from 3 cultures. Mean ± s.e.m. One-way ANOVA with Dunnett’s multiple test correction. **c.** Top, experimental design. Bottom, representative IF images of puromycin incorporation in DRG neurons. *Ahr*^cKO^ neurons exhibited significantly higher labeling than controls. Average intensity of n = 20 randomly selected fields of independent DRG neuron cultures from 2 mice per group. Mean ± s.e.m. Unpaired two-tailed Student’s t-test. **d.** Top, experimental scheme of in vivo puromycin incorporation assay. Bottom, IF images and quantification showing increased neuronal puromycin labeling in cKO DRGs compared to controls. Quantification of mean fluorescence intensity of puromycin from n = 3 L5 DRG collected from 3 mice per genotype. Mean ± s.e.m. Unpaired two-tailed Student’s t-test. **e.** Heatmap of key translation regulators identified as AhR-responsive by RNA-seq of ipsi- and contralateral DRGs at 1 dpi after PL (n = 5 per condition; adjusted *P* value < 0.05). **f.** Diagram of translational regulators affected by *Ahr* cKO in response to PL. **g.** IF showing elevated p-RPS6 (Ser235/236) in cKO DRG neurons compared to controls at 1 dpi, with minimal signal in glia. Quantification of mean p-RPS6 fluorescence intensity from randomly selected fields of sciatic DRGs of n = 5 control and n = 4 cKO mice. Mean ± s.e.m. Unpaired two-tailed Student’s t-test.

When comparing Puro incorporation in primary DRG cultures at 24 h post-seeding (note that axotomy occurs during dissociation), we observed that *Ahr*^cKO^ neurons on average displayed a 25% higher Puro incorporation than control cells (**Fig. 6c**). Similarly, in vivo puromycin injection at 1 dpi after PL confirmed significantly greater puro labeling (24% higher) in *Ahr*^cKO^ DRG neurons (**Fig. 6d**). Thus, AhR-deficient DRG neurons displayed enhances de novo protein synthesis after axotomy.

To further investigate underlying mechanisms, analysis of the DRG RNA-seq data for translational regulators revealed that *Ahr*^cKO^ DRGs showed reduced expression of *Eif4ebp1* (an inhibitor of 5′ cap-dependent translation)*, Sesn* and *Castor* (amino acid sensors and inhibitors of mTOR activity), along with decreased *Rps14*, encoding a 40S ribosomal subunit protein (**Fig. 6e, f**). In line with these changes, phosphorylated riboprotein S6 (p-RPS6^Ser235/236^), a marker of mTOR activity, was elevated in *Ahr*^cKO^ DRG neurons (**Fig. 6g**). Together, these data indicate that AhR constrains neuronal translation by limiting mTOR signaling and ribosome processing, whereas its loss unleashes a neuron-intrinsic increase in protein synthesis to support regenerative growth.

### Axon growth-promoting effect of AhR ablation requires HIF-1α

Because AhR and HIF-1α share ARNT as a dimerization partner, we next tested whether HIF activity is required for the regenerative phenotype of *Ahr*^cKO^ neurons (**Fig. 7a**). Pharmacological inhibition of HIF translation with KC7F2 ^54^ reduced HIF abundance and abolished the growth advantage of *Ahr*^cKO^ neurons (**Fig. 7b**; **Fig. S11a**). Likewise, siRNA-mediated *Arnt* knockdown also eliminated the axon-promoting effect of *Ahr* deletion, reducing neurite length to control levels (**Fig. 7c**), indicating that the HIF-ARNT pathway is engaged during axon regrowth of *Ahr*^cKO^ DRG neurons.

**Figure 7.**
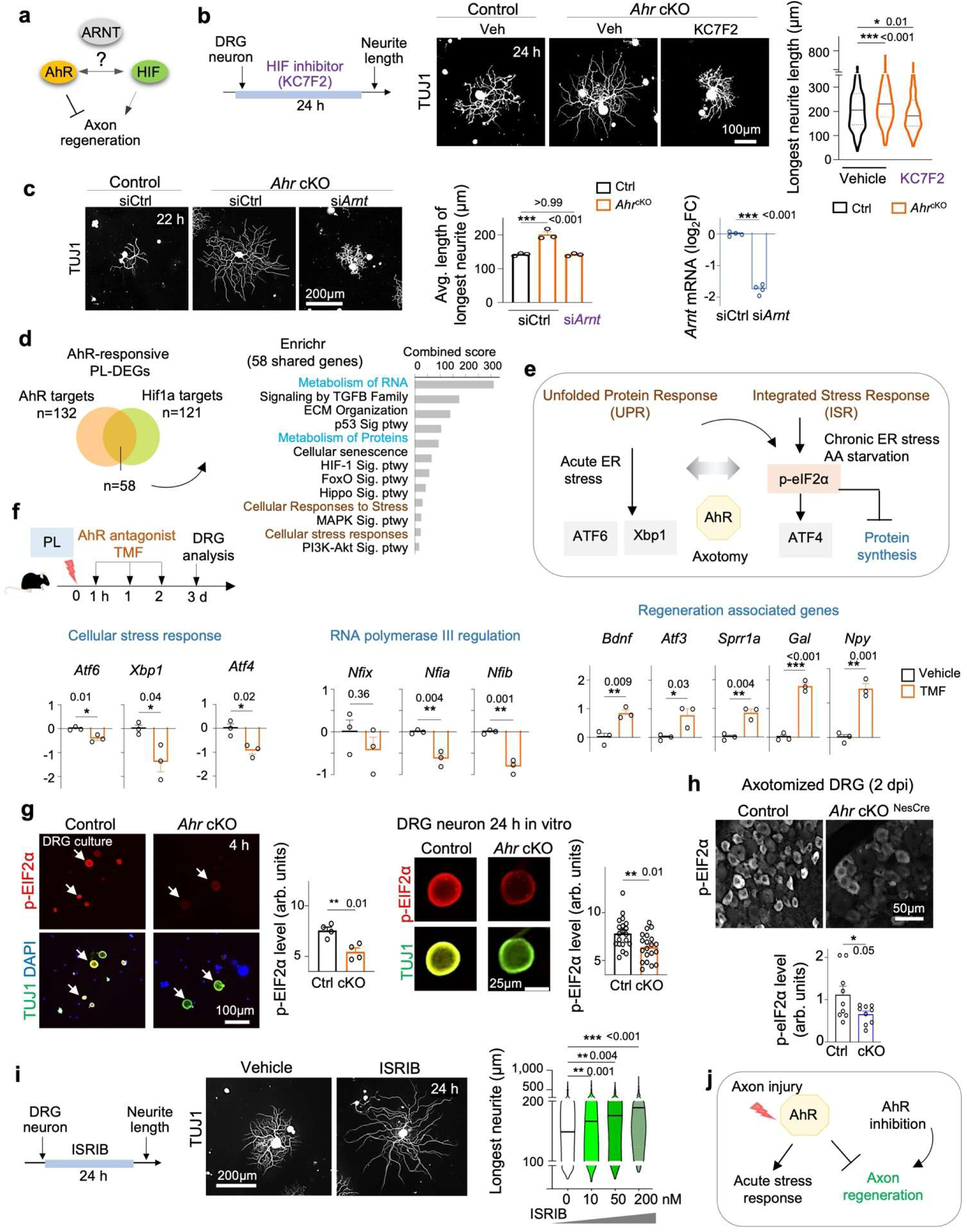
Growth-promoting effect of *Ahr* deletion requires crosstalk with HIF1α. **a.** Schematic of AhR-HIF interactions in regulating axon regeneration. **b.** Left, experimental scheme of HIF inhibitor KC7F2 (10 μM). Right, IF images and quantification show that KC7F2 abolished the neurite outgrowth advantage of *Ahr*^cKO^ DRG neurons. Violin plots show median and quartiles. n = 250-408 neurons per condition, from independent cultures of 2 mice per genotype. One-way ANOVA with Dunnett’s correction. **c.** Left, IF and quantification show that siRNA knockdown of *Arnt* reverted the enhanced neurite growth of *Ahr*^cKO^ neurons. Quantification of mean longest neurite length from n = 10 fields across 3 independent cultures per genotype. One-way ANOVA with Dunnett’s correction. Right, qRT-PCR confirms reduced Arnt mRNA 48 h after siRNA treatment. n = 4 per group. Unpaired two-tailed Student’s *t*-test. Mean ± s.e.m. **d.** Venn diagram of overlapping genes (n = 58) between putative AhR and HIF targets, with pathway enrichment analysis (right). **e.** Diagram of stress sensors linked to protein synthesis regulation. **f.** Top, scheme of post-injury administration of AhR antagonist TMF (10 mg/kg, i.p., daily × 3) or vehicle after sciatic nerve crush; DRGs collected at 3 dpi. Bottom, qRT-PCR normalized to *Hprt1* shows TMF suppressed stress-associated genes and Pol III regulators while inducing regeneration-associated genes. n = 3 independent samples from 3 mice, each pooled from L4-L6 DRGs. One-way ANOVA with Dunnett’s correction. Mean ± s.e.m. **g.** IF images show reduced p-eIF2α (Ser51) in *Ahr*^cKO^ neurons versus controls at 4 h and 24 h in vitro. Quantification of mean fluorescence intensity: n = 4 fields at 4 h, n = 20 fields at 24 h per genotype. Unpaired two-tailed Student’s t-test. Mean ± s.e.m. **h.** Reduced p-eIF2α levels in DRG neurons in vivo at 2 dpi after PL in *Ahr*^cKO^ mice versus controls. Quantification from sciatic DRGs of n = 3 mice per genotype. Unpaired two-tailed Student’s t-test. Mean ± s.e.m. **i.** Left, scheme. Right, dose-dependent effect of ISR inhibitor (ISRIB) on neurite outgrowth. n = 625 (0 nM), 462 (10 nM), 474 (50 nM), 427 (200 nM) neurons from 4 cultures. One-way ANOVA with Dunnett’s correction. Mean ± s.e.m. **j.** Working model: AhR induction after axonal injury promotes acute stress responses and proteostasis, constraining growth; AhR inhibition relieves these restraints to enable axon regeneration.

In contrast, conditional deletion of *Arnt* in vivo (Thy1-CreERT2/EYFP *Arnt*^fl/fl^) did not impair baseline or PL-induced axon outgrowth in primary DRG neurons, and regenerating SCG10^+^ axons were comparable between *Arnt* mutants and controls at 3 dpi (**Fig. S11b-e**). Thus, while *Ahr*^cKO^ neurons require ARNT for their enhanced growth, ARNT-deficient neurons can compensate via alternative pathways (**Fig. S11f**).

To further investigate the interplay of neuronal AhR with the HIF pathway, we applied ChEA3 analysis to the AhR-responsive PL-DEGs, which identified 121 candidate HIF targets, nearly half overlapping with putative AhR targets **(Fig. 7d**). The 58 overlapping genes were enriched for RNA and protein metabolism, cellular stress responses, cellular senescence, and axon growth promoting signaling such as HIF-1, p53, MAPK, PI3K-Akt (**Fig. 7d**; **Table S6**). These findings support transcriptional crosstalk between HIF and AhR after axotomy; AhR inhibition shifts the balance from AhR-mediated cellular stress responses and proteostasis towards HIF-mediated metabolic adaption and pro-growth signaling.

Integration with epigenomic data after PL implicated 5hmC-mediated regulation in the AhR-HIF crosstalk. Our previous genome-wide profiling of PL-induced differentially hydroxymethylated regions (DhMRs) identified 1,036 sites residing mostly at gene bodies (rather than promoters) ^3,55^, and they were significantly enriched for motifs of bHLH-PAS TFs including HIF-1α, ARNT, Bmal1, but not AhR (**Fig. S12a, b**). Notably, 40% of DhMRs contained HIF motifs, corresponding to 766 genes, with 27 overlapping with AhR-responsive PL-DEGs, and they were enriched for amino sugar metabolism, ubiquitination, neuronal migration, and Wnt signaling (**Fig. S12c, d**; **Table S7**). Thus, AhR-responsive PL-DEGs may operate in part through DhMR/HRE in *Ahr*^cKO^ DRG.

Further linking AhR and 5hmC, *Ahr*^cKO^ DRGs showed elevated 5hmC levels after axotomy (**Fig. S12e**). In our prior study, we found that circadian regulator Bmal1 gates regenerative responses through 5hmC^3^. Interestingly, xenobiotic metabolism and AhR signaling were among the top enriched pathways in the Bmal1 cKO regulon, implicating Bmal1 as a potential upstream regulator of AhR signaling (**Fig. S12f**). Together, these data place AhR, HIF-1α, and Bmal1 as interconnected BHLH-PAS TFs that orchestrate transcriptional and epigenomic programs in response to axonal injury (**Fig. S12g**).

### AhR mediates cellular stress response after axotomy

To define putative neuronal AhR regulon, we leveraged published scRNA-seq of DRG neurons across 6 h to 14 dpi after PL ^56^, which identified 98 AhR-associated genes (**Fig. S13a, b**; **Table S8**). Unlike regeneration-associated TFs (ATF3, Sox11, Smad1, Creb1, Myc, Jun) or early response genes (Egr1, Fos), AhR was predicted to be inactivated during axon regeneration, in line with its transient activation and an inhibitory role of AhR on axon regeneration. Echoing our RNA-seq results of *Ahr*^cKO^ DRGs, pathway analysis of the putative neuronal AhR regulon (n=98 genes) highlighted translational control (e.g., RNA Pol III transcriptional termination, regulation of transcription by RNA Pol II, Negative regulation of pri-miRNA, and RNA Pol III transcriptional termination) (**Fig. S13c**). Indeed, the AhR regulon included *Nfia* and *Nfib,* both nuclear factors involved in RNA Pol III transcriptional termination, and both carry AHREs in their promoters (**Fig. S13d**). The involvement of RNA Pol III implicates small RNA biogenesis (tRNA, 5S rRNA^57^) in AhR-mediated translational control of neuronal genes.

Notably, neuronal AhR regulon also included stress-response TFs ATF4 and ATF6 (**Fig. S13b**). ATF4 is a sensor of integrated stress response (ISR) in response to chronic ER stress, amino acid starvation, or viral infection, etc. ^58^, and it is activated by increased cellular levels of uncharged tRNA ^59^ (**Fig. 7e**). scRNA-seq showed that both *Atf4* and *Atf6* were induced in most DRG neurons, and both contained AHREs in their promoters, and this also applied to *Xbp1*, another TF regulating acute ER stress (**Fig. 7e**; **Fig. S14a, b**). Validation studies showed that post-injury administration of pharmacological inhibitors of AhR in vivo suppressed stress-responsive TFs (*Atf4, Atf6*, *Xbp1)* and RNA Pol III regulators *(Nfia, Nfib*), while upregulating generation-associated genes (RAGs; *Bdnf, Atf3, Sprr1a, Gal, Npy*), as well as *Stmn1* (encoding SCG10)*, Tead1* (Hippo signaling), and other AHRE/HRE-bearing genes such as *Sox9, Rest, Cdk6* (**Fig. 7f**; **Fig. S14c, d**).

Phosphorylation of eIF2α (p-eIF2α^Ser51^) is a central integration step of unfolded protein response (UPR) and ISR that leads to inhibition of protein synthesis ^58^ (**Fig. 7e**). We examined p-eIF2α in DRG neurons and found that *Ahr*^cKO^ neurons exhibited reduced p-eIF2α levels (a sign of increased protein translation), both in vitro (4 h and 24 h post-seeding) and in vivo after peripheral axotomy (**Fig. 7g, h**). In functional assays, blocking the ISR with the small molecule compound ISRIB ^60^ significantly enhanced neurite outgrowth of DRG neurons in a dose-dependent manner (**Fig. 7i**). Together, these findings support the model that AhR induction after axotomy leads to proteostasis and translational suppression via ISR/UPR, whereas inhibition of AhR can relieve this brake and redirect neuronal responses toward metabolic adaption and pro-regrowth signaling (**Fig. 7j**).

## DISCUSSION

Neurons must balance environmental stressors with regenerative demands after axonal injury. Here, we identify AhR as a central regulator of this stress-growth switch. The induction of AhR after axotomy engages proteostasis and integrated stress response (ISR) to preserve tissue integrity but constrains regeneration; AhR inhibition shifts the balance toward elevated translation and metabolism, promoting axon regrowth in both PNS and CNS injury models.

AhR is known as a xenobiotic sensor that regulates detoxification enzymes and transporters. Interestingly, evolutionary evidence suggests that detoxification is a later adaptation of a more ancestral role in neurons ^61^. As such, AhR is broadly expressed in the nervous system ^61,62^, regulates axon and dendrite branching in invertebrates ^62,63^, and in mammals, its loss enhances dendritic complexity and neuronal differentiation ^64,65^. Our findings extend this paradigm, establishing AhR as a neuronal brake on axon regeneration, in line with computations from unbiased high-throughput screens predicting AhR as a regulator of axon growth ^66^.

Mechanistically, AhR induction after axotomy rewires specific gene programs that reinforce protein quality control and proteostasis. AhR activation after PL was transient, dampened by multiple feedback loops ^6^. This transience ensures an acute stress response, while freeing ARNT for HIF signaling in a timely fashion. Indeed, in our axon regrowth studies, *Ahr*^cKO^ neurons required HIF-ARNT for enhanced growth, with shared transcriptional targets enriched in metabolic and regenerative pathways.

AhR engaged broad proteostasis programs, spanning RNA Pol III activity, ribosome assembly, autophagy, ubiquitination, protein trafficking, and mitochondrial-cytosolic crosstalk, whereas its loss activated growth-promoting pathways (HIF, BDNF, PI3K-Akt) and micronutrient metabolism. The AhR injury regulon also featured stress-associated TFs (ATF4, ATF6, Xbp1), and AhR-deficient neurons showed increased protein synthesis, elevated p-RPS6, and reduced p-eIF2α, consistent with relief from translational suppression. Hence, the AhR regulon enforces proteostasis and UPR/ISR programs, while constraining axon regeneration. We further uncovered crosstalk among bHLH-PAS transcription factors. Circadian factor Bmal1 appears to be upstream of xenobiotic pathways, while HIF also regulates circadian programs ^67^. Integration with epigenomic datasets revealed that AhR-responsive PL-DEGs overlap with PL-induced DhMRs harboring HIF motifs. AhR cKO increased 5hmC marks in DRG neurons after injury, suggesting transcriptome rewiring through epigenetic remodeling. Together, AhR, HIF, and Bmal1 form an interconnected network coordinating stress adaptation and regeneration.

Functionally, AhR cKO enhanced axon regrowth and motor-sensory recovery after both PNS (peripheral nerve) and CNS (spinal cord) lesions. Post-injury treatment with the AhR antagonist SR1 phenocopied AhR cKO effects, although the promoting effect was limited in the CNS paradigm due to relatively robust regeneration capacity of PNS neurons. As SR1 affects hematopoietic stem cells ^68^, additional studies are needed to define drug specificity, target engagement, pharmacokinetics, dosing, timing of AhR modulators and side effects. The identity of endogenous AhR ligands remains elusive, as these may potentially arise from local metabolites, systemic inflammation, or oxidative stress ^69^. We found that antibiotics-mediated microbiome depletion had no overt effect on nerve regeneration in our experimental paradigms, though other gut related effects on AhR activity cannot be excluded. Interestingly, constitutive Cyp1a1 expression can partially phenocopy AhR loss by depleting endogenous ligands ^9^, thus suggesting alternative routes of AhR modulation. Furthermore, AhR may also act on gene expression via non-AHRE interactions ^70^.

More broadly, our results align with protective effects of AhR inhibition observed in several CNS disorders, including stroke ^71^, Huntington’s disease ^72^, and gut-lung ^73^ or endothelial responses to environmental cues ^74,75^. Natural AhR polymorphisms shape ligand sensitivity across species ^6^, raising the possibility that regenerative capacity varies with diet, pathogens, or genetics.

In summary, our studies establish AhR as a brake on axon regeneration that integrates transcriptional, metabolic, and epigenetic programs to enforce proteostasis at the expense of regenerative growth. By placing AhR within a regulatory network together with HIF and Bmal1, our work opens door for alternative strategies to target stress-growth plasticity for improved nervous system repair.

## METHODS

### Animals

All animal studies were performed under protocols approved by the Institutional Animal Care and Use Committee (IACUC) at Icahn School of Medicine at Mount Sinai and the Institutional Review Board/Animal Ethics Committee of Texas A&M University. Animals were housed in a pathogen-free barrier facility, under a 12-hour light:dark cycle with food and water provided ad libitum.

C57BL/6J mice (JAX stock no. 000664), *Ahr*^fl/fl^ mice (*Ahr*^tm3.1Bra^/J; stock no. 006203), Nestin-Cre mice (B6.Cg-Tg(Nes-cre)1Kln/J; stock no. 003771), Thy1-CreER^T2^ mice (Tg(Thy1-cre/ERT2,-EYFP)HGfng/PyngJ; stock no. 012708), and Rosa26^INTACT^ mice (B6;129-Gt(ROSA)26Sor^tm5(CAG-Sun1/sfGFP)Nat^/J; stock no. 021039) were obtained from Jackson Laboratory. *Arnt*^fl/fl^ mice were obtained from Frank Gonzalez, NIH. All mice were bred on a C57BL/6J genetic background. Littermates were used as controls.

The following primers were used for genotyping by PCR using mouse tail DNA:

*Ahr*^fl/fl^ mice: F1: GTCACTCAGCATTACACTTTCTA, F2: CAGTGGGAATAAGGCAAGAGTGA, R1: GGTACAAGTGCACATGCCTGC. Expected band sizes: 106 bp for wildtype allele, 140 bp for floxed allele, and 180 bp for excised floxed allele.

*Arnt*^fl/fl^ mice: F1: TGCCAACATGTGCCACCATGT, R1: GTGAGGCAGATTTCTTCCATGCTC. 290 bp for wild type allele, 340 bp for Arnt floxed allele.

Nestin-Cre mice: F1: CCGCTTCCGCTGGGTCACTGT, R1: TGAGCAGCTGGTTCTGCTCCT, R2: ACCGGCAAACGGACAGAAGCA. 379 bp for wild type allele, 229 bp for transgenic Cre allele. Rosa26^INTACT^ mice: F1: GCACTTGCTCTCCCAAAGTC, R1: CATAGTCTAACTCGCGACACTG, R2: GTTATGTAACGCGGAACTCC. 557 bp for wild type allele, 300 bp for knockin allele.

Thy1-CreER^T2^ mice: F1: TCTGAGTGGCAAAGGACCTTAGG, R1: CGCTGAACTTGTGGCCGTTTACG, Int-F2: CAAATGTTGCTTGTCTGGTG, Int-R2: GTCAGTCGAGTGCACAGTTT. 200 bp for wild type allele, 300 bp for transgenic Cre allele.

### Pharmacological treatments

To induce CreER-mediated conditional knockouts, tamoxifen (Sigma T5648) in corn oil (Sigma C8267) was injected into adult mice intraperitoneally (100 mg/kg) once daily for 5 days.

AhR agonists: 2-(1′H-indole-3′-carbonyl)-thiazole-4-carboxylic acid methyl ester (ITE; Tocris 1803), L-kynurenine (L-Kyn; Tocris 4393), 6-formylindolo[3,2-b]carbazole (FICZ; Tocris 5304), and norisoboldine (NOR; Selleckchem S9092) were reconstituted in DMSO. For in vitro studies, applied concentrations are indicated in the Results section. For in vivo experiments, ITE was diluted in 12.5% kolliphor/PBS (Sigma C5135) to a final volume of 600 µl and injected intraperitoneally at 10 mg/kg.

AhR antagonists: CH-223191 (CH; Tocris 3858), 6,2′,4′-trimethoxyflavone (TMF; Tocris 3859), StemRegenin-1 (SR1; Selleckchem S2858), and BAY 2416964 (Selleckchem S8995) were reconstituted in DMSO. For in vivo studies, TMF was further diluted in 12.5% kolliphor/PBS. TMF was injected intraperitoneally at 10 mg/kg, and SR1 and BAY 2416964 at 25 mg/kg using a Hamilton fine syringe (Hamilton 80920).

HIF translational inhibition: To inhibit HIF translation, KC7F2 (Cayman 14123), a small-molecule inhibitor targeting HIF via translational control was used.

Integrated stress response (ISR) inhibition: ISRIB (Sigma-Aldrich 5.09584) was reconstituted in DMSO. Antibiotics treatment: Broad-spectrum depletion of gut microbes was performed following a published protocol with minor modifications ^47^. Briefly, wild-type C57BL/6J mice received drinking water containing ampicillin sodium (1 g/l; Sigma A9518), vancomycin hydrochloride (0.5 g/l; Sigma V2002), neomycin sulfate (1 g/l; Sigma N1876), metronidazole (1 g/l; Sigma M1547), sucrose (50 g/l; Sigma S0389), and acetic acid (4 mM) for 18 days prior to sciatic nerve crush. Solutions were provided ad libitum in light-protected bottles and replaced every third day.

Indole metabolite: 3-Indolepropionic acid (IPA; Selleckchem S4809) was dissolved in DMSO.

As vehicle controls for drug treatments, either DMSO or DMSO dissolved in 12.5% kolliphor/PBS were used as appropriate.

### Sciatic nerve lesion

Male and female mice, age 8-18 weeks old unless otherwise specified, were anesthetized by isoflurane inhalation (5% for induction, 2% for maintenance). A small skin incision was made at mid-thigh using a scalpel blade after skin preparation and disinfection. To clearly expose the sciatic nerve, the fascial space between biceps femoris and gluteus superficialis muscles was opened gently without causing hemorrhage. The nerve was then freed from surrounding connective tissue under microscope, avoiding shearing or traction forces. For sciatic nerve crush model, the nerve was crushed with an Ultra-fine Hemostat (Fine Science Tools 13020-12) for 15 sec at the third click. For sciatic nerve transection model, the nerve was cut with a 3 mm Vannas Spring Scissor (Fine Science Tools 15000-00). For sham surgery, the sciatic nerve was exposed as described but left intact. Mouse skin was closed with Reflex 7 mm Wound Closure System (Braintree Scientific RF7 CS) after surgery. Mice were left to recover in a warm cage. All surgical instruments were autoclaved before surgery and principles of asepsis were maintained throughout.

### Spinal cord injury model

T7-T9 laminectomy was performed on 8-week-old mice (wild-type C57Bl/6 female). The mice were then clamped using 2 pairs of Adson forceps before using the Infinite Horizon Impactor at 60 kdyn of force with a 2 sec dwell time to induce a moderate T8 contusion and compression as described ^76–78^. The muscles and skin were secured with 5.0 sutures, and the skin was sealed with Dermabond. After surgery, mice recovered in a warmed cage and then moved to a normally temperate cage and provided with food and water ad libitum. All animals received subcutaneous injections of 1 ml saline, 10 mg/kg Baytril, and 0.05 mg/kg buprenorphine daily for the first week post-surgery. All surgeries were performed in the morning (8am - 12pm) to limit potential circadian influence. Bladders were expressed manually twice daily. All drug administrations were done by individuals blinded to experimental groups.

### DRG isolation

DRG dissections were conducted under a Nikon SMZ645 stereomicroscope. Mice were euthanized by carbon dioxide inhalation followed by cervical dislocation. Animals were positioned supine and secured to a dissection pad. The ventral thoracic and abdominal skin and viscera were removed to expose the ventral spinal column using surgical scissors (Fine Science Tools, 14054-13) with tissue forceps (Fine Science Tools, 11021-12) for support. Ventral paraspinal muscles were cleared using spring scissors (Fine Science Tools, 15751-11) to visualize the lumbosacral peripheral nerves. To expose the lumbosacral DRGs, ventral vertebral elements were removed with the same spring scissors, aided by octagon forceps (Fine Science Tools, 11042-08), taking care to avoid nerve transection. L4-6 DRGs were then isolated by gently elevating the ganglion with Dumont #3 forceps (World Precision Instruments, 50037) and severing connecting nerves with Vannas spring scissors (Fine Science Tools, 15000-00).

### Primary DRG neuron culture

Adult DRGs from adult C57BL/6J mice were dissected and placed in ice-cold DMEM/F12 (Gibco 11330057). DRGs were washed 3 times with ice-cold calcium- and magnesium-free HBSS (Gibco 14175095) including 10 mM HEPES (Gibco 15630106) before incubating in 0.3% collagenase I (Worthington LS004196) for 90 min at 37 °C. DRGs were then washed 3 times with HBSS buffer with HEPES at room temperature, followed by additional digestion in 0.25% trypsin-EDTA (Gibco 25200072) containing 50 μg/ml DNase I (Worthington LK003172) for 30 min at 37 °C. Trypsinization was stopped with warm DMEM medium (Gibco 10569044) containing 10% FBS (Gibco 26140079) and DNase I.

DRGs were dissociated by trituration with fire-polished Pasteur glass pipets (Fisherbrand 13-678-20D). To remove myelin debris and cell clumps, a partial-purification step was performed by centrifugation through a BSA (Fisher BP9700100) cushion. Specifically, the DRG suspension was mixed with 8 ml NS-A Basal Medium (NeuroCult 05750) and then 2 ml of 15% BSA in HBSS was added at the bottom of the 15 ml centrifuge tube followed by centrifugation at 1,000 rpm for 6 min. Supernatant was carefully removed and DRGs were resuspended in NS-A Basal medium containing 2% B27 (Gibco A3582801), 0.725% glucose, 0.5 mM L-glutamine, and 0.4% antibiotic-antimycotic (Gibco 15240062). DRG neurons were plated on poly-L-ornithine (Sigma P4957) and laminin (Gibco 23017015) coated chamber slides or 6-well plates for subsequent experiments. siRNA-mediated knockdown studies were conducted as previously described with modification ^79^. ∼4,000 DRG neurons were resuspended in 1.5 ml of titration media (without DNase I) and gently mixed with 0.5 ml of transfection complex containing 2 μl of DharmaFECT 2.0 (Dharmacon, #T-2002-02) and 2 μl of siRNA at 20 μM stock concentration in Neurocult NB-A media and seeded on PLO/Laminin coated plates/coverslips. ON-TARGETplus SMART pool siRNA oligos were obtained from Dharmacon (Ahr siRNA-L-044066-00, Arnt siRNA-L-040639-01-0005, and non-targeting pool, D-001810-10-05).

### Generation of induced human neurons

The studies using hESCs were approved by the Embryonic Stem Cell Research Oversight Committee (ESCRO) at Icahn School of Medicine at Mount Sinai. Induced neurons were generated as described ^3^. Briefly, H9 ESCs were induced to neuroprogenitor cells (NPCs) using STEMDiff SMADi neural induction kit (Stem Cell Technologies 08581) as instructed. NPCs were passaged at a density of 1.2 ×10^6^ in 2 ml of STEMdiff Neural Progenitor Medium (Stem Cell Technologies 05833). Mixed cortical neuron culture was induced from NPCs as described ^80^. Differentiation media contained BrainPhys media (StemCell Technologies 05790) with 1x N2 (Gibco 17502048), 1x B27 (Invitrogen 12587-010), 20 ng/ml brain-derived neurotrophic factor (BDNF, Peprotech 450-02), 20 ng/ml glia-derived neurotrophic factor (GDNF, Peprotech 450-10), 200 µM L-ascorbic acid (Stem cell technologies 72132), and 250 µg/ml dibutyryl cyclic AMP sodium salt (db-cAMP, Stem Cell Technologies 73884). Half the media volume was changed with fresh differentiation media every other day for 10 to 13 days before analysis.

### Mouse cortical neuron culture

Cortical adult neuron assay was conducted as described ^33^. In brief, wild-type 6-week-old C57BL/6 male mice were euthanized using CO_2_ and the cortex was isolated and transferred to a MACS C-tube, then dissociated using the Miltenyi gentleMACS octo-dissociator on a preset protocol. This was followed by the removal of debris and endothelial blood cells using the Mitlenyi MACS Adult Brain Dissociation Kit (Miltenyi Biotec 130-107-677). Next, using the Adult Neuron Isolation Kit (Miltenyi Biotec 130-126-603), following manufacturer’s instructions, the negative fraction (fraction enriched with neurons) was collected and used for the cortical neurite outgrowth assay.

For cultures of early postnatal cortical neurons, cortices from P0–P2 WT mouse pups were isolated after careful removal of meninges. Tissue was minced, washed in cold HBSS, and dissociated using the Neural Tissue Dissociation Kit-T (Miltenyi, #130-094-802). Following cell counting, 10 × 10^4^ cells were seeded per well onto 24-well plates containing glass coverslips coated with poly-L-ornithine (Sigma, #P4957) and laminin (Gibco, #23017015).

### Neurite outgrowth assays

L4-L6 DRG neurons were seeded on pre-coated 4-well chamber slides (Falcon 10384501) at ∼1,000-2,000 neurons per well. Neurons were fixed with ice-cold 4% PFA and stained with anti-Tubulin β3 (TUJ1) to visualize outgrowing neurites.

For pharmacological experiments, DRG neurons from wild-type C57BL/6J mice were plated on pre-a coated 6-well plate, using neurons from 8-10 DRGs per well. Neurons were cultured for 24 h with pharmacological AhR modulators. Cells were either fixed for immunofluorescence staining or used directly for RNA lysis and qRT-PCR analysis.

A replating assay was performed on induced neurons between differentiation day 10 to 13 as described ^3,81^. Briefly, cells were washed twice with PBS and incubated in 0.025% trypsin for 5 min at 37 °C. Trypsin was gentle removed while keeping neurons attached and replaced with differentiation media. Gentle pipetting was then carried out to dissociate the neurons followed by counting and seeding in 4-well chamber slides at a density of 55,000 cells/well in differentiation media containing Ahr agonists or antagonists for 1 day before analysis.

Adult mouse cortical neurite outgrowth assay was performed as described ^33^. Briefly, primary adult cortical neurons were seeded onto PDL coated plastic bottom plates (Greiner-Bio 781091) at 10,000 cells/well for 2 days. StemRegenin1 (SR1, TargetMol T1831) and vehicle were added at the time of plating and left in the media for 2 days without media change. Neuronal media consisted of MACS Neuro Media (Miltenyi Biotec 130-093-570), 2 mM L-alanine-L-glutamine dipeptide (Sigma-Aldrich G8541), and 1x B27 Plus supplement (ThermoFisher A3582801).

For postnatal mouse cortical neurite outgrowth assays, at seeding, cultures were treated with Ahr modulators or vehicle control in Neurobasal-A medium (Gibco, #10888022) supplemented with 2% B27 Plus (Gibco, #A3582801), 2 mM GlutaMAX-I (Gibco, #35050061), 5% FBS, and 1% penicillin-streptomycin (Gibco, #15140122), and maintained at 37 °C with 5% CO_2_. Neurons were fixed after 24 hours for immunostaining, imaging, and quantification of of neurite length.

### Puromycylation (SUnSET) Assay for Nascent Protein Synthesis

Puromycin dihydrochloride (MP Biomedicals, 210055280; Sigma-Aldrich, P8833) was dissolved in water to generate a 10 mg/ml stock solution. For optimization of the use in neuronal cultures, puromycin was applied at 0.3 - 10 µg/ml for up to 30 min at 37 °C, followed by three washes with ice-cold PBS and immunofluorescence analysis using an anti-puromycin antibody. Vehicle–treated cultures served as negative controls and were processed in parallel. For labeling of DRG neurons, cultures were pulsed with 1 µg/ml puromycin in fresh medium for 10 min at 37 °C, washed twice with ice-cold PBS, and analyzed as above.

In vivo puromycylation assay was performed as previously described ^82,83^. Puromycin was prepared as a 4.8 mg/ml stock in PBS and administered intraperitoneally at 21.8 mg/kg (0.040 µmol/g). Mice were euthanized 30 min post-injection; DRGs were dissected, fixed in 4% PFA for 1.5 h at 4 °C, and processed for immunofluorescence with anti-puromycin antibody.

### RNA isolation and qRT-PCR

Total RNA of cells or tissues was extracted with RNeasy Plus Mini kit (QIAGEN 74134). For RNA collection from tissue, dissected DRGs were initially stored in RNAlater Stabilization Solution (Invitrogen AM7024) and then homogenized in RLT Plus buffer including 1% β-mercaptoehanol using RNase-free disposable pellet pestles (Fisherbrand 12-141-364). For RNA collection from cells, cell cultures were washed once with PBS and then lysed by vigorous pipetting. Genomic DNA was eliminated through a gDNA eliminator column according to the manufacturer’s instructions. RNA was eluted in RNase-free water and stored at −80°C. cDNA was prepared with the SuperScript III First-Strand Synthesis System (Invitrogen 18080051) from equal amounts of RNA (approximately 200 ng from DRG tissues and 500 ng from cell culture for each reaction). Quantitative RT-PCR (qRT-PCR) was performed with PerfeCTa SYBR Green FastMix Rox (Quanta Bioscience 95073-012) with an ABI 7900HT quantitative PCR system (Applied Biosystems) at the Mount Sinai qPCR CoRE. *Hprt1* was used as the house-keeping gene to normalize qRT-PCR results. Primers for qRT-PCR analysis are listed in **Table S9**.

### Western blot

L4-L6 DRG were collected and immediately frozen in liquid nitrogen and stored at −80 °C for later analysis. Tissues were homogenized and lysed with RIPA buffer (Sigma R0278) containing EDTA-free protease inhibitor cocktail (Roche 04693159001) and phosSTOP (Roche 4906845001). The frozen DRG in a 1.5 ml tube were disrupted with RNase-free disposable pellet pestles (Fisherbrand 12-141-364) on ice. 1 U/μl benzonase nuclease (Millipore E1014) was added to lysis buffer. Samples were mixed on a rotator at 4 °C for 30 min and then spun in a tabletop centrifuge at 13,000 rpm for 10 min to remove undissolved pellet. An equal volume of 4x LDS sample buffer (Invitrogen NP0008) was added to lysates, which were then boiled at 95 °C for 5 min. Lysates from equal number of DRGs were loaded and separated by electrophoresis on 4-12% ExpressPlus gels (Genscript M41210), followed by transfer to a PVDF membrane. Membranes were blocked in Intercept Blocking Buffer (LI-COR Biosciences 927-70001) at room temperature for 1 h and subsequently incubated with primary antibodies diluted with Intercept Antibody Diluent (LI-COR Biosciences 927-75001) at 4 °C overnight. Blots were washed with PBST (5 x 5 min) and incubated with secondary antibodies at room temperature for 1 h. Bands were detected with the Odyssey Infrared Imaging System (LI-COR Biosciences) and band intensity was quantified with Image Studio 5.2.5 (LI-COR Biosciences).

Primary antibodies for Western blots:

anti-AHR (rabbit, used as 1:1,000 dilution, Enzo BML-SA210),

anti-CYP1B1 (rabbit, 1:1,000, Invitrogen PA5-95277),

anti-HIF1α (rabbit, 1:500, Novus NB100-479),

anti-ATF3 (rabbit, 1:1,000, Santa Cruz sc-188),

anti-β-Actin (mouse, 1:10,000, Sigma A1978).

Secondary antibodies for Western blots:

800CW donkey anti-rabbit IgG (1:10,000, LI-COR Biosciences 926-32213),

680RD donkey anti-mouse IgG (1:10,000, LI-COR Biosciences 926-68072).

### Immunofluorescence

For immunofluorescence (IF) of cultured cells, cultures were washed once with PBS and then fixed in 4% ice-cold PFA for 15 min. For IF of cryosections of DRG and sciatic nerves, tissues were fixed in 4% ice-cold PFA/PBS for 12 hr, washed in PBS, soaked in 30% sucrose overnight, and then embedded in OCT compound (Fisher Scientific 23-730-571). Cryosections were cut at 12 μm thickness and placed on SuperFrost Plus slides (VWR 48311-703) and stored at −20 °C before analysis. Sections were washed with PBS and incubated in blocking buffer containing 5% normal donkey serum (Jackson Immunoresearch 017-000-121) and 0.3% Triton X-100 (Acros Organics 9002-93-1) in PBS for 1 h at room temperature. Primary antibodies were diluted in antibody dilution buffer containing 1% BSA (Fisher BioReagents BP9700100) and 0.3% Triton X-100 in PBS and incubated at 4 °C overnight. Alexa-coupled secondary antibodies were diluted in antibody dilution buffer and added on sections after three washes with PBS and incubated for 1 h at room temperature. DAPI (Invitrogen D1306) was used for nuclear counterstaining (1:1,000). Slides were washed thrice with PBS and mounted with Fluromount G (Southern Biotech 0100-01). Wholemount staining of footpad was performed as previously described ^3^. Briefly, the footpad skin of the injured hind paw was dissected, cleaned from connective tissue, washed with PBS, and fixed in 4% PFA overnight at 4 °C. Tissue was rinsed 10x for 30 min with PBS containing 0.3% Triton-X (0.3% PBST) followed by incubation with primary antibody in blocking buffer (0.3% PBST containing 5% goat serum and 20% DMSO) for 5 days at room-temperature (RT) with gentle shaking. Tissue was washed 10x for 30 min with 0.3% PBST and incubated with secondary antibody in blocking buffer for 3 days at RT with gentle shaking. Subsequently, tissue was washed 10x for 30 min with 0.3% PBST, dehydrated in 50% methanol for 5 min, 100% methanol for 20 min, and cleared in a 1:2 benzyl alcohol: benzyl benzoate mix overnight at room temperature. Imaging was performed in clearance solution with a Zeiss LSM 780 confocal microscope.

Primary antibodies for IF:

anti-AHR (rabbit, 1:300, Enzo BML-SA210),

anti-ATF3 (rabbit, 1:300, Santa Cruz sc-188),

anti-Tubulin β3 (TUJ1, mouse, 1:1,000, Biolegend 801201),

anti-Tubulin β3 (D71G9, rabbit, 1:300, Cell Signaling 5568S),

anti-SCG10/STMN2 (rabbit, 1:1,000, Novus NBP1-49461),

anti-GFP (chicken, 1:1,000, Aves Lab GFP-1020),

anti-IBA1 (rabbit, 1:1,000, Wako 019-19741),

anti-pS6 ribosomal protein-S235/236 (rabbit, 1:300, Cell Signaling 2211),

anti-5hmC (rabbit, 1:500, Active Motif 39769),

anti-CD8a (rat, 1:100, Invitrogen 14-0081-82),

anti-CD4 (rat, 1:100, Invitrogen 14-0041-82),

anti-CD68 (rat, 1:100, Biolegend 137002),

anti-CD45 (rat, 1:100, BD Pharmingen 550539),

anti-PU1 (rabbit, 1:300, Thermofisher MA5-15064),

anti-F4/80 (rat, 1:300, Thermofisher 14-4801-81),

anti-CD206 (goat, 1:200, R&D systems AF2535),

anti-Sox10 (goat, 1:50, R&D systems AF2864),

anti-PGP9.5 (rabbit, 1:800, Neuomics, RA12103),

anti-NFH (chicken, EMD Millipore AB5539, 1:1,000),

anti-CSPG (CS-56) (mouse, Sigma C8035, 1:100),

anti-GFAP (chicken, Aves Labs, 1:500),

anti-puromycin (mouse, DSHB, PMY-2A4, 1:100),

anti-phospho-Eif2a (S51) (rabbit, Cell Signaling, 3398, 1:300),

anti-E-Cadherin (24E10) (rabbit, Cell Signaling 3195, 1:300),

anti-Ly6G (rat, Biolegend 127601, 1:100),

anti-Hif1a (rabbit, Novus NB100-479, 1:300).

Alexa-conjugated donkey secondary antibodies (Jackson ImmunoResearch) were used at 1:300 dilution of a 1 mg/ml stock solution (in 50% glycerol):

AlexaFluor 488 anti-rabbit IgG (711-545-152),

AlexaFluor 488 anti-chicken IgY (703-545-155),

AlexaFluor 594 anti-rabbit IgG (711-585-152),

AlexaFluor 594 anti-mouse IgG (711-585-150),

AlexaFluor 594 anti-rat IgG (712-585-153),

AlexaFluor 647 anti-rabbit IgG (711-605-152),

AlexaFluor 647 anti-mouse IgG (715-605-151).

### Mouse intestinal tissue preparation

Mouse intestines were prepared according to the Swiss-roll methodology that allows efficient analysis of epithelial morphology ^84,85^. Briefly, mouse intestines were isolated from freshly euthanized mice and placed immediately in ice cold PBS. Intestines were carefully handled with forceps and flushed multiple times using a 20 ml syringe to clear stool and any remaining debris. At this stage, the colon and small intestines were cut and handled separately. A 1 ml glass pipette was inserted into the tissue and carefully laid on a large piece of Whatman filter paper. A sharp blade was used to cut intestines longitudinally down the length of the pipette which was then lightly rolled sideways to flatten the tissue on filter paper. The flattened tissue was subsequently rolled on a Gmark cotton swab stick and immersed in ice cold 4% PFA for overnight fixation at 4 °C. Next day, intestines were washed 3x with ice cold PBS and soaked in 15% sucrose/PBS followed by 30% sucrose/PBS solution each overnight to preserve tissue morphology. Tissues were subsequently embedded in OCT, sectioned, and stained as described above.

### Motor and sensory behavioral testing

All behavioral tests were performed by an investigator blinded to the cohort identities. All animals were acclimated to the isolated procedure room for 30 min before testing. For motor function recovery testing after sciatic nerve injury, hindpaw prints were collected before and after sciatic nerve crush injury. Hindpaws were pressed on an ink pad and mice were then allowed to walk on white paper to collect the prints. The Sciatic Functional Index (SFI) was calculated by measuring dimensions of the paw prints ^86^, using the following formula: SFI = −38.3x(EPL-NPL)/NPL + 109.5x(ETS-NTS)/NTS + 13.3x(EIT-NIT)/NIT - 8.8 (E-, experimental-; N-, normal-; PL, print pength; TS, total spread; IT, intermediate toes). For analysis of sensory functional recovery, von Frey filament tests were performed ^87^. The plantar surface of the hindpaw was pricked with a series of fine filaments and the mechanical threshold that evoked a withdrawal reflex was recorded. For ladder walking test, regular rungs were spaced evenly at 1 cm and irregular rungs were arranged in a pseudo-random pattern with variable spacing (1–3 cm). Mice were allowed to walk the length of the ladder while a video recorded. Each hindlimb step was categorized as correct placement, partial slip, or full slip. The number of errors was normalized to the total number of steps to calculate an error rate for each animal. To conduct open field Basso Mouse Scale (BMS) testing, mice were placed in an open field for 5 minutes to allow two observers to evaluate freely roaming mice by assessing the following parameters: ankle movements, stepping pattern, coordination, paw placement, trunk stability, and tail movement. The mice were scored according to the BMS scoring system ^76,88,89^. Mice were tested prior to injury (baseline), 2 days post injury, and then weekly thereafter. Mice with BMS scores above 5 at 2 days post injury (incomplete injuries) or any mouse that died prematurely before the end of the study were excluded from analysis.

### RNA-seq analysis

Contralateral and ipsilateral DRGs were collected from control or Ahr cKO mice at 1 dpi by sciatic nerve transection. For next-generation sequencing, RNA was isolated from DRG tissue with the QIAGEN RNeasy plus mini kit, and cDNA libraries were sequenced on the Illumina NovaSeq platform (Psomagen). Preprocessing, quality control, and alignment of FASTQ files was performed using the NGS-Data-Charmer pipeline. Trim-Galore tool (v0.6.5) ^90^ was used for adaptor trimming and alignment to the mouse mm10 genome assembly was performed with Bowtie2 (v2.4.1) ^91^. The ‘rmdup’ module of SAMtools (v1.10) ^92^ was used to remove duplicated read pairs. FeatureCounts was used to obtain a gene expression matrix, using the parameters “--fraction -t gene” on the GENCODE annotation (vM25). Genes with fewer than 5 samples showing a minimum read count of 2 reads were filtered out before performing differential gene expression analysis with DESeq2. For visualization of genome-wide RNA-seq read distribution, aligned BAM files were further processed into BigWig file format and visualized in the Integrative Genomics Viewer (IGV)^93^ for inspection of *Ahr* exon read coverage.

### Image analysis

Fluorescence images of mouse DRG neurons, human induced neurons, sciatic nerve, DRG, and intestinal tissues were captured using a Zeiss Axioscope microscope with AxioCam MRm camera and quantifications were performed using Image J (version 2.3.0/1.53q) as previously detailed ^3^.

For mouse adult cortical neuron culture, images were acquired using ImageXpress (IXM) Micro Confocal High-Content Imaging System (Molecular Devices). The length of the longest neurite of each neurite-bearing neuron (neurite longer than the diameter of its soma) was measured using the Simple Neurite Tracer (SNT version 4.0.3) plugin. The percentage of neurite bearing neurons was calculated by counting neurons with neurites longer than the diameter of soma relative to total neurons. Quantification of cytoplasmic to nuclear shuttling was performed by measuring nuclear signal relative to total signal for each individual neuron for both cultured cells and DRGs. For DRG tissue image analysis, the threshold function was used to quantify percent area per section or cell number relative to total determined by DAPI staining. Quantification of cell markers in intestines was conducted by manual cell counting in multiple villi per section.

Adult mouse cortical neuron images were quantified using the Neurite Outgrowth Analysis Module in MetaXpress 6 software (Molecular Devices). The number of valid neurons was determined by quantifying the number of TUBB3^+^/DAPI^+^ cells in a well with ≥ 10 µm of total neurite outgrowth. Total neurite outgrowth was determined by dividing the length of all neurites in a well by the number of valid neurons in that respective well.

To establish a regeneration index of injured sciatic nerves, tiled images were merged using Photoshop CC 2019. The SCG10 fluorescence intensity was measured along the length of the nerve using ImageJ. A rectangular region-of-interest (ROI) containing the lesion site and adjacent proximal and distal areas was selected to generate a ‘plot profile’. The position with maximal SCG10 profile intensity was used to normalize the regeneration index and the position with minimal intensity was used for subtraction of background value. Most distal SCG10 fluorescence intensity above background was used to determine maximal axonal length. For skin reinnervation analysis, maximal intensity projections of 400 μm z-stack images were used for quantification of percentage of PGP9.5^+^ puncta normalized to the area of footpad using image J.

### Bioinformatics

TF interaction networks were generated using STRING database ^21^, with the default setting of medium confidence. Ahr and Hif1a putative target genes were identified using ChIP-X Enrichment Analysis with ChEA3 ^53^. Heatmaps, volcano plots, bubble plots, bar graphs, box plots, and violin plots were generated with FLASKi ^94^, OriginLab, and Graphpad Prism 10.4.2. Gene set enrichment analysis (GSEA) was performed using GSEA_4.3.2 software provided by the Broad Institute ^95^, using the non-preranked whole DRG genome and Hallmark_MSigDB gene sets. Pathway enrichment in gene sets was performed with Enrichr ^96,97^ and IPA Qiagen knowledge database ^98^ with the whole list of expressed genes as background. Identification of experimentally validated promoter motifs was conducted using the Eukaryotic Promoter Database platform ^99^.

### Statistical analysis

For each dataset, Shapiro-Wilk test was performed to test data normality (*P* > 0.05 determined as parametric and *P* < 0.05 determined as nonparametric). For parametric data, unpaired two-tailed Student’s *t*-test was used for comparison between two groups, one-way analysis of variance (ANOVA) with Holm-Sidak multiple test correction for comparison of three groups, and Two-way ANOVA followed by Bonferroni’s multiple comparisons test for multiple groups For nonparametric data, Mann-Whitney two-tailed *t*-test was used for comparison between two groups and Kruskal-Wallis test with Dunn’s multiple test correction for comparison between multiple groups. All statistical analyses were performed with GraphPad Prism version 9 or 10. The GraphPad Prism setting NEJM (New England Journal of Medicine) for reporting of *P* values was applied. *P* ≤ 0.05 was considered as statistically significant (*); *P* ≤ 0.01, **; *P* ≤ 0.001, ***.

## Data Availability

Source data are provided with this paper. The *Ahr* cKO RNA-seq datasets are deposited at the NCBI GEO database under accession numbers GSE243308 and GSE307639.

## Acknowledgements

We are grateful to all members from the Zou laboratory for help and suggestions. We thank Frank Gonzalez, NIH, for providing the Arnt-flox mouse line.

This work was supported by grants of the NIH (R01 NS127442), the New York State SCIRB (Spinal Cord Injury Research Board) (DOH01-C33268GG, DOH01-C30832GG, and DOH01-C32242GG), and the Neilsen Foundation (#890112) to H.Z, and by NIH R21NS131834 and R01NS128101 grants to C.G.G.. Y.W. was partly supported by a scholarship fund from Xi’an Jiaotong University.

## Author contributions

Conceptualization: DHalaw, YW, CCG, RHF, HZ; Data acquisition: DHalaw, YW, JL, DHalp, HN, AS; Data analysis: DHalaw, YW, AS, CCG, RHF, HZ; Bioinformatic analysis: ME, DHalaw, YW, AR, LS; Manuscript writing: DHalaw, YW, RHF, HZ.

