## Supplementary figures for "AhR restricts axon regeneration by balancing neuronal stress and growth response after injury"

**Supplementary Figures 1-14**

**Supplementary Tables**

**Table S1.** PL-DEGs of DRGs (ipsi/contra)

**Table S2.** PL-DEGs of AhR cKO DRGs (ipsi/contra)

**Table S3.** Venn diagram gene lists

**Table S4**. Ahr responsive xenobiotic metabolism genes (n=134)

**Table S5.** Shifted genes in AhR cKO after PL

**Table S6.** Putative AhR (n = 132), HIF (n=121), and overlap target genes with transcriptional shift in AhR cKO after PL

**Table S7.** Venn diagram of gene lists

**Table S8.** AhR neuronal regulon (n = 98) from scRNA analysis

**Table S9.** qRT-PCR primers

**Supplementary Videos: Functional recovery after spinal cord injury (SCI)**

**Video S1 & S2.** Control mouse (video S1) and *Ahr* cKO mouse (video S2) recorded in a regular rung ladder test at 35 dpi after T8 contusion SCI.

**Video S3 & S4.** Control mouse (video S3) and *Ahr* cKO mouse (video S4) recorded in an irregular rung ladder test at 35 dpi after T8 contusion SCI.

**Video S5 & S6. C**ontrol mouse (video S5) and an *Ahr* cKO mouse (video S6) recorded in open field performance to score for hindlimb motor function with the Basso Mouse Scale (BMS) after T8 contusion SCI.

**Video S7 & S8.** Vehicle-treated control mouse (video S7) and SR1-treated mouse (video S8) recorded in a regular rung ladder test at 35 dpi after T8 contusion SCI.

**Video S9 & S10.** Vehicle-treated control mouse (video S9) and SR1-treated mouse (video S10) recorded in a regular rung ladder test at 35 dpi after T8 contusion SCI.

**Video S11 & S12.** Vehicle-treated control mouse (video S11) and SR1-treated mouse (video S12) recorded in open field performance to score for hindlimb motor function with BMS after T8 contusion SCI.

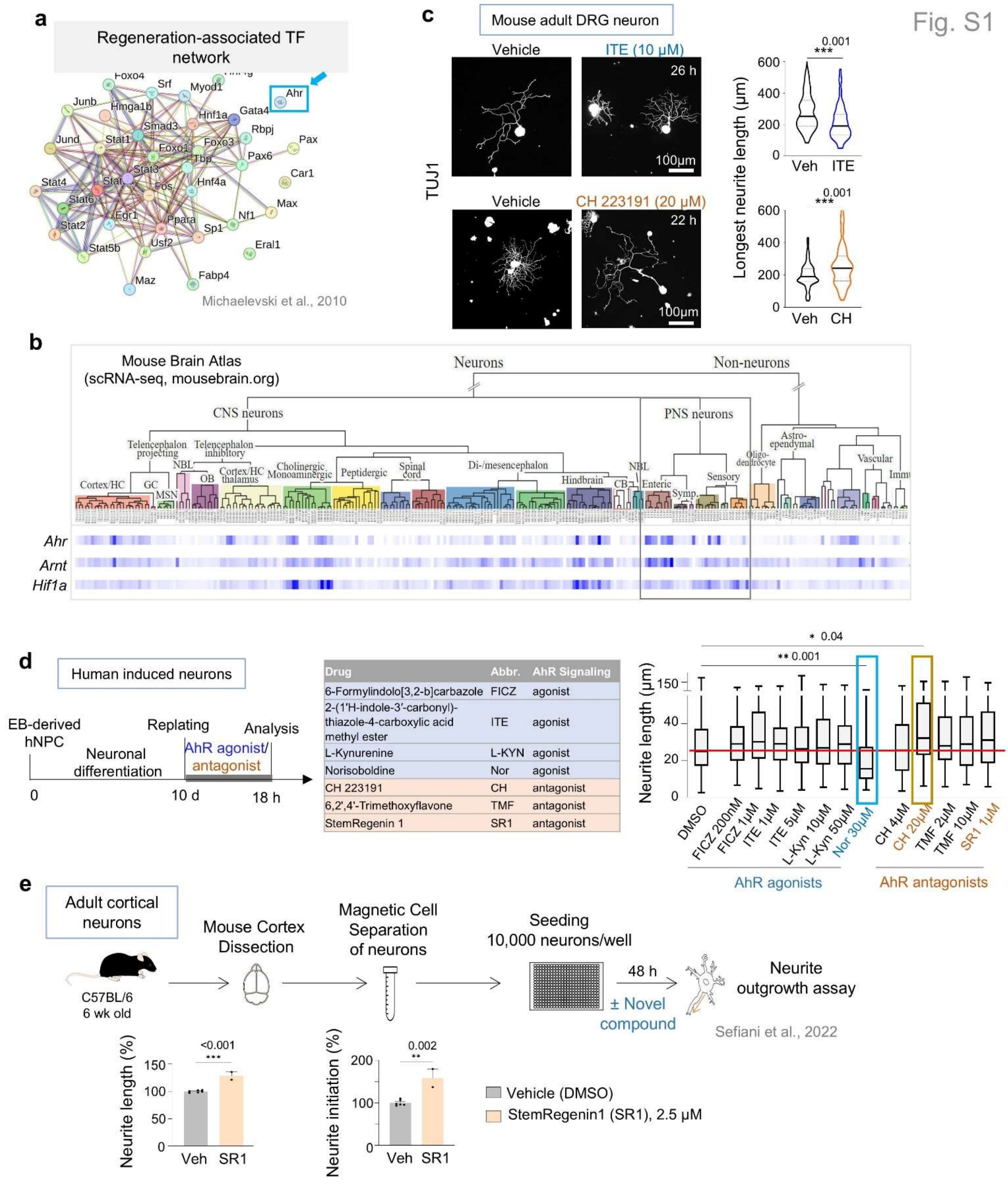


| Drug | Abbr. | AhR Signaling |
|---|---|---|
| 6-Formylindolo[3,2-b]carbazole | FICZ | agonist |
| 2-(1'H-indole-3'-carbonyl)-thiazole-4-carboxylic acid methyl ester | ITE | agonist |
| L-Kynurenine | L-KYN | agonist |
| Norisoboldine | Nor | agonist |
| CH 223191 | CH | antagonist |
| 6,2',4'-Trimethoxyflavone | TMF | antagonist |
| StemRegenin 1 | SR1 | antagonist |

**Figure S1. AhR inhibitor enhances neurite outgrowth.**

**a.** STRING protein-protein interaction network of transcription factors (TFs) induced after conditioning lesion of DRG, identified in Michaelevski et al., 2010.

**b.** Heatmap of *Ahr, Arnt,* and *Hif1a* expression across cell types of the mouse nervous system (scRNA-seq atlas; Zeisel et al., 2018).

**c.** IF images and quantification of TUJ1+ neurites in adult DRG neurons treated with AhR agonist ITE (10 μM), antagonist CH-223191 (20 μM), or vehicle for one day post-plating. ITE study: n = 74 neurons (vehicle), 100 (ITE), from 2 mice. CH study: n = 120 neurons (vehicle), 188 (CH), from 2 mice. Median and quartiles shown. Mann-Whitney two-tailed test.

**d.** Left, schematic of AhR agonist/antagonist stimulation of induced neurons derived from hESCs. Middle, table of AhR pharmacological modulators. Right, box plots of neurite length after 18 hr of treatment (n = 70-116 neurons). Median and quartiles shown. Kruskal-Wallis test with Dunn's correction; only significant *P* values shown. Highlighted conditions (Nor, CH, SR1) correspond to main Fig. 2.

**e.** Top, schematic of high-throughput screen to identify regeneration-promoting compounds in adult cortical neurons (Sefiani et al., 2022). Bottom, quantification of effects of AhR antagonist SR1 on total neurite length per neuron and neurite initiation. Each data point represents mean of >150 neurons (with neurites ≥ 10 μm) from a single well. n = 2-6 wells per group. Mean ± s.e.m. Unpaired two-tailed Student's t-test.

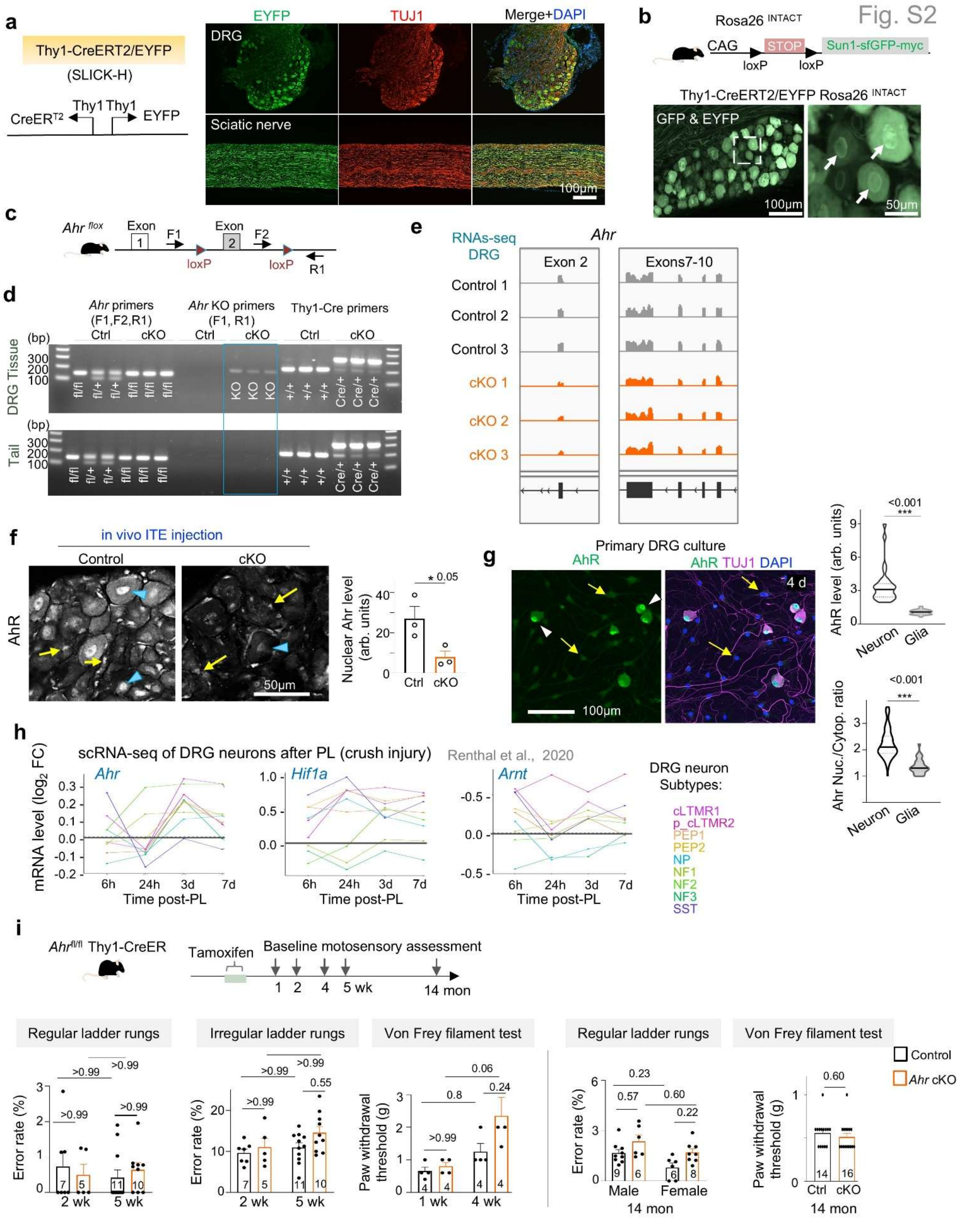


**Figure S2. Validation of neuronal *Ahr* cKO mice and baseline motosensory performance.**

**a.** Diagram of the SLICK-H transgenic line expressing tamoxifen-inducible CreERT2 and EYFP under bidirectional Thy1 promoter. IF images of DRG and sciatic nerve from Thy1-CreERT2/EYFP mice show pan-neuronal ($TUJ1^+$) EYFP expression (IF with anti-GFP antibody). DAPI, nuclear counterstain.

**b.** Top, schematic of $Rosa26^{INTACT}$ (INTACT, isolation of nuclei tagged in specific cell types) reporter allele (Mo et al., 2015). Bottom, IF of DRG section from $Rosa26^{INTACT}$ Thy1-CreERT2/EYFP mouse shows expression of cytoplasmic GFP from Thy1-CreERT2/EYFP allele and nuclear membrane GFP from INTACT/Sun1-GFP reporter (arrows).

**c.** Schematic of the floxed *Ahr* allele with loxP sites flanking exon 2 and position of genotyping primers.

**d.** PCR genotyping of *Ahr*$^{fl/fl}$ Thy1-CreERT2/EYFP mice after tamoxifen (100 mg/kg i.p. x 5 d; analysis 14 d later) confirms excision of *Ahr* exon 2 in DRG but not tail DNA. Expected bands: 106 bp (WT), 140 bp (floxed), 180 bp (excised).

**e.** RNA-seq read tracks (3 examples for each genotype) show reduced *Ahr* exon 2, but no change in downstream exons 7 - 10 in DRGs of *Ahr*$^{cKO}$ mice as compared to DRGs in control mice (contralateral DRGs of sciatic nerve injury study were used).

**f.** IF of DRGs from mice treated in vivo with ITE show high nuclear levels of AhR in neurons, which was ablated in *Ahr*$^{cKO}$ DRGs (blue arrowheads; contralateral DRGs used). No change was observed in glial AhR levels (yellow arrows), which displayed lower levels than neurons. n = 3 L5 DRGs per condition. Mean ± s.e.m. Unpaired two-tailed Student's t-test.

**g.** IF of primary DRG cultures shows higher nuclear AhR signals in $TUJ1^+$ neurons (arrowheads) compared to glia (yellow arrows). Quantification: AhR intensity, n = 30 cells; nuclear/cytoplasmic AhR ratio, n = 100 neurons and 30 glia from 3 cultures. Violin plots show median and quartiles. Unpaired two-tailed Student's t-test.

**h.** Expression of *Ahr, Hif1a*, and *Arnt* across DRG neuron subtypes after PL, generated from scRNA-seq data of study (Renthal et al., 2012).

**i.** Top, experimental timeline of assessment of motosensory performance after neuronal *Ahr* cKO at baseline without injury. Bottom, quantifications at different timepoints after induction of cKO. Sample sizes are indicated in graphs. Mean ± s.e.m. Two-way ANOVA with Bonferroni correction; Mann-Whitney test for von Frey assay.

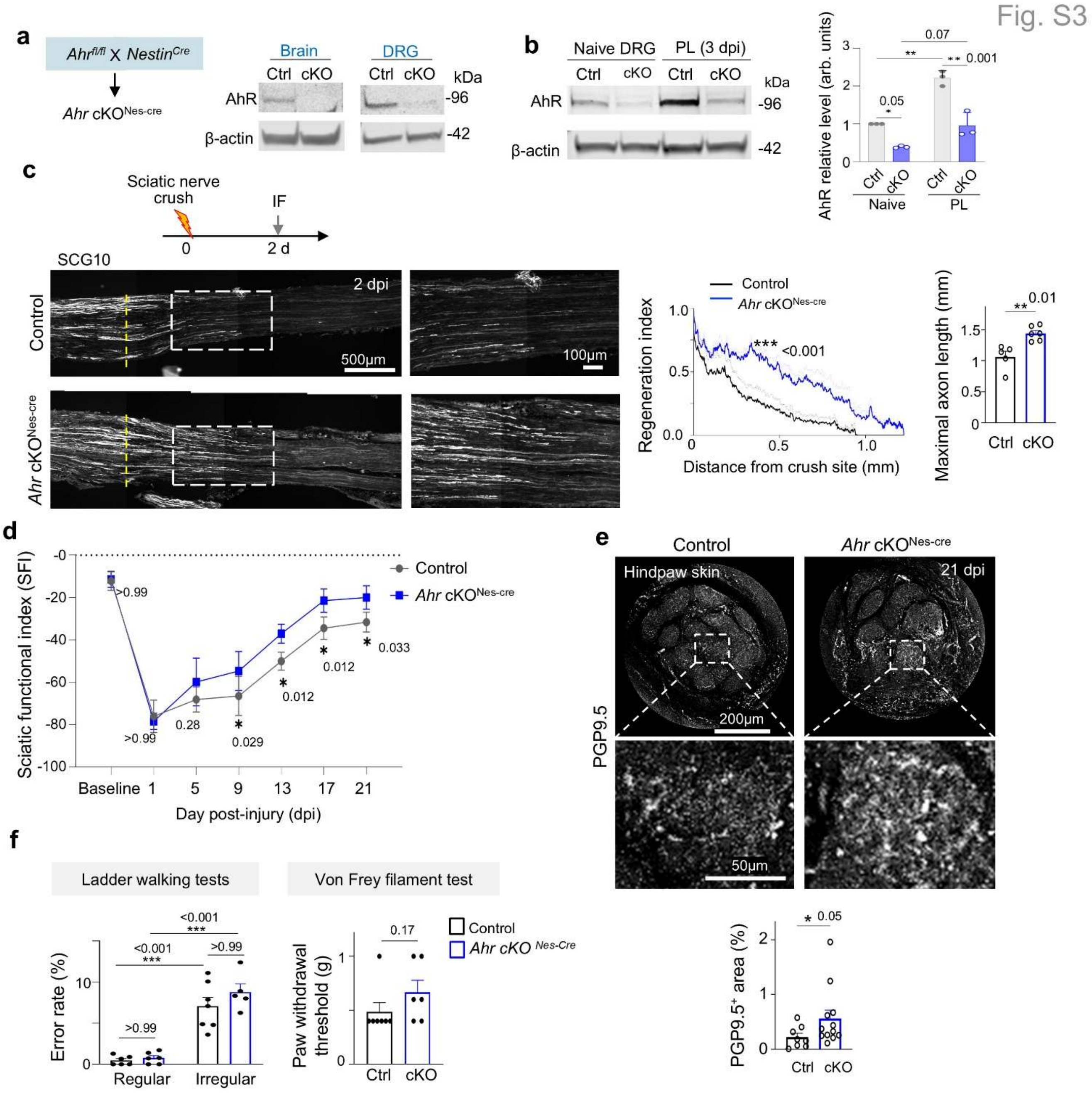


**Figure S3. *Ahr* cKO (Nestin-Cre) accelerates axon regeneration after sciatic nerve injury.**

**a.** Schematic of the *Ahr*$^{cKO}$ (Nestin-Cre) allele. Immunoblots show marked reduction of AhR protein in brain and DRG from this allele. β-actin, loading control.

**b.** Immunoblots and quantification of AhR protein in naive and conditioned DRG (3 dpi after PL) from *Ahr*$^{cKO}$ (Nestin-Cre) and control mice. n = 3 samples (each from pooled L4-L6 DRGs) from 3 mice per genotype. Mean ± s.e.m. Two-way ANOVA with Bonferroni correction.

**c.** Top, experimental paradigm. Bottom, IF images of SCG10$^{+}$ axons traversing lesion center (dashed vertical line) at 2 dpi in *Ahr*$^{cKO}$ (Nestin-Cre) and controls. Right, quantification of regeneration index and maximal axon length. n = 5 control, n = 6 cKO mice. Mean ± s.e.m. Regeneration index: two-way ANOVA with Bonferroni correction. Maximal axon length: unpaired two-tailed Student's t-test.

**d.** Sciatic functional index (SFI) measurements show improved motor recovery in cKO mice. n = 5 per group. Mean ± s.e.m. Two-way ANOVA with Bonferroni correction.

**e.** Whole-mount IF for PGP9.5 shows enhanced epidermal re-innervation in cKO mice at 21 dpi. Images represent maximal-intensity projections of ~400 μm footpad sections spanning top, middle, and lower epidermis. Quantification of PGP9.5$^+$ area from two footpads per mouse. n = 4 control, n = 6 cKO mice. Mean ± s.e.m. Unpaired two-tailed Student's t-test.

**f.** Baseline motosensory tests. n = 7 control, n = 6 cKO mice. Mean ± s.e.m. Ladder walking: two-way ANOVA with Bonferroni correction. Von Frey assay: Mann-Whitney two-tailed test.

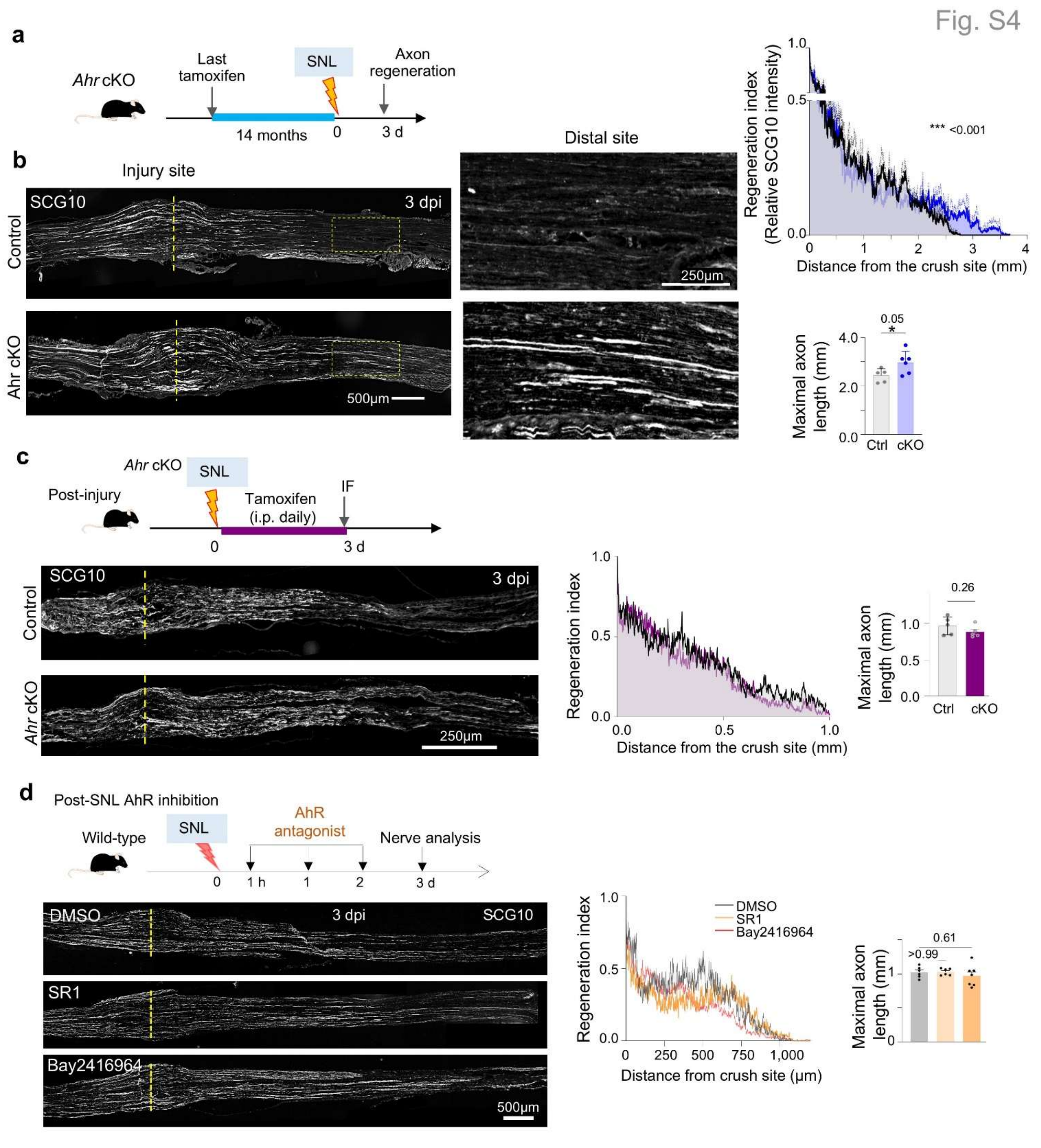


**Figure S4. Neuronal *Ahr* deletion enhances peripheral nerve regeneration in older mice, but AhR inhibition post-injury has no effect.**

**a.** Long-term neuronal *Ahr* cKO paradigm.

**b.** Tamoxifen (100 mg/kg i.p., daily x 5) was administered 14 months before sciatic nerve crush. At 3 dpi, IF of SCG10$^{+}$ axons showed enhanced regeneration in cKO mice compared to controls. Quantification of regeneration index (two-way ANOVA with Bonferroni correction) and maximal axon length distal to lesion center (n = 5 control, n = 6 cKO; unpaired two-tailed Student's t-test). Mean ± s.e.m.

**c.** Post-injury *Ahr* cKO paradigm. Tamoxifen (100 mg/kg i.p., daily x 3) was administered immediately after sciatic nerve crush. At 3 dpi, $SCG10^{+}$ axon regeneration was comparable between cKO and control mice (n = 5 per group). Quantification of regeneration index and maximal axon length distal to lesion center shown (unpaired two-tailed Student's t-test). Mean ± s.e.m.

**d.** Post-injury AhR antagonist treatment. SR1 (25 mg/kg), Bay2416964 (25 mg/kg), or vehicle (DMSO) were administered daily x 3 immediately after sciatic nerve crush. At 3 dpi, IF of $SCG10^{+}$ axons showed no significant differences between groups. n = 6 mice (DMSO, SR1), n = 7 (Bay). Mean ± s.e.m. One-way ANOVA with Dunnett's correction.

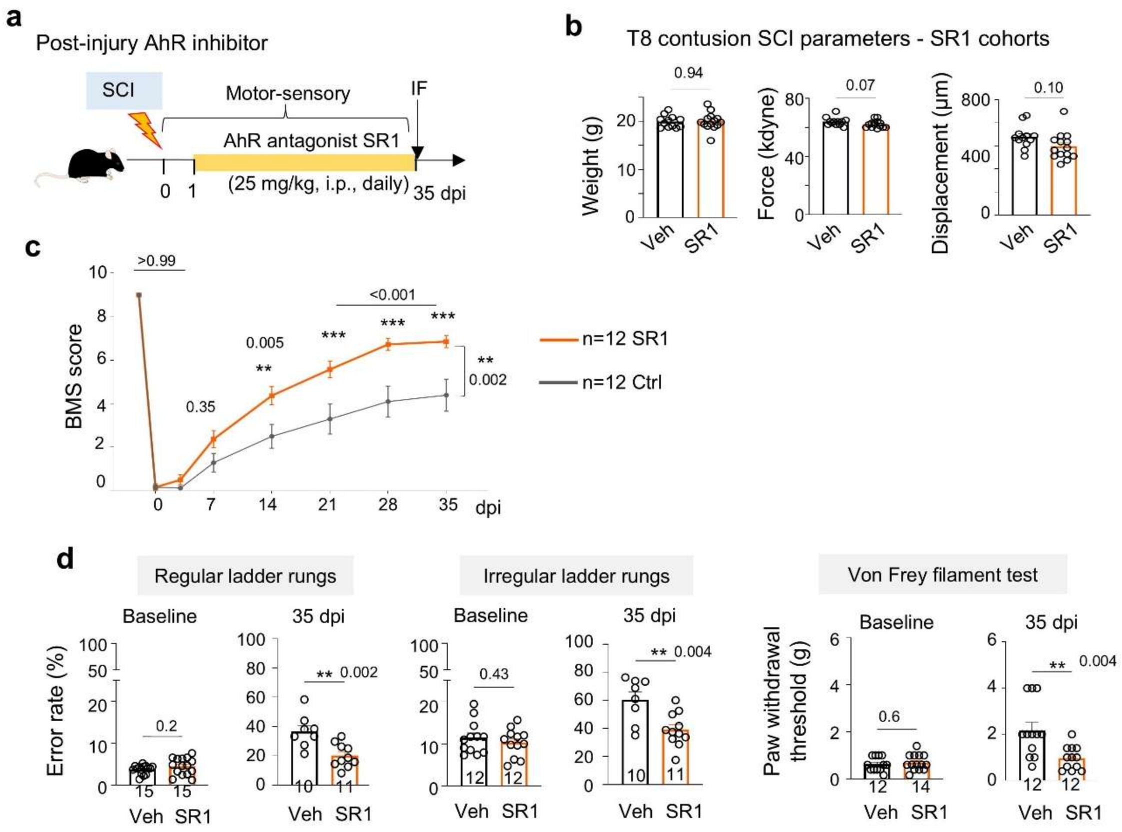


**Figure S5. Post-injury AhR inhibition enhances functional recovery after spinal cord injury.**

**a.** Experimental scheme for in vivo administration of AhR antagonist SR1 starting at 1 day post-SCI, (25 mg/kg i.p., daily).

**b.** Similar parameters of contusion injury with impactor device in SR1 and control cohorts. Mean ± s.e.m. unpaired two-tailed Student's *t*-test. n = 12 control and n=14 mice per group

**c.** Basso Mouse Scale (BMS) locomotor scores show improved recovery in mice treated with SR1 after SCI compared to controls. n = 12 mice per group. Mean ± s.e.m. Two-way ANOVA with Bonferroni correction.

**d.** Ladder walking (regular and irregular rungs) and von Frey filament testing were performed at baseline and 35 dpi. Post-SCI treatment with SR1 improved motor coordination and sensory recovery. Mouse cohort size indicated in the graph. Data represent mean ± s.e.m.; unpaired two-tailed Student's t-test.

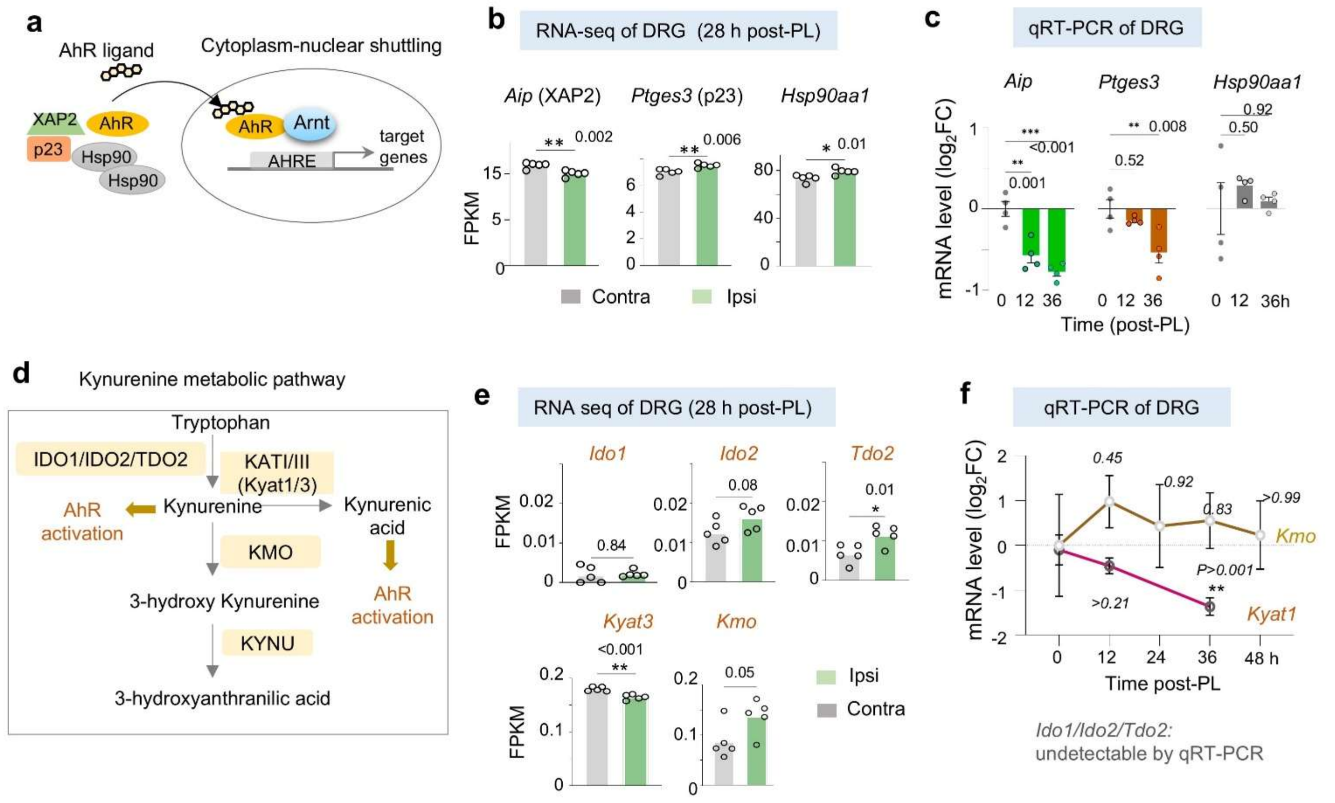


**Figure S6. Expression of Kynurenine pathway genes after PL.**

**a.** Diagram of the cytoplasmic AhR complex. AhR is sequestered in cytoplasm by two Hsp90 chaperones, co-chaperone p23 (*Ptges3*), and XAP2 (*Aip*). Ligand binding triggers dissociation of the complex and nuclear translocation.

**b.** RNA-seq results for *Aip, Ptges3,* and *Hsp90* expression in ipsi and contralateral sciatic DRGs (FPKM, fragements per kb transcript per million reads). n = 5 samples per condition. Mean ± s.e.m. Unpaired two-tailed Student's t-test.

**c.** Time-course qRT-PCR analysis of *Aip, Ptges3,* and *Hsp90aa1* expression at 12 h and 36 h post-PL. n = 4 samples per condition. Mean ± s.e.m. One-way ANOVA with Dunnett's correction.

**d.** Overview of tryptophan metabolism / kynurenine pathway: IDO, indoleamine 2,3 dioxygenase; TDO, tryptophan-2,3-dioxygenase; KYAT1, kynurenine amino transferase1; KMO, kynurenine 3 monooxygenase; KYNU, kynureninase.

**e.** RNA-seq data of kynurenine pathway genes in ipsi- and contralateral sciatic DRGs. n = 5 per condition. Mean ± s.e.m. Unpaired two-tailed Student's t-test.

**f.** Time-course qRT-PCR of *Kmo* and *Kyat1* expression in axotomized DRG. n = 3 per condition. Mean ± s.e.m. One-way ANOVA with Dunnett's correction.

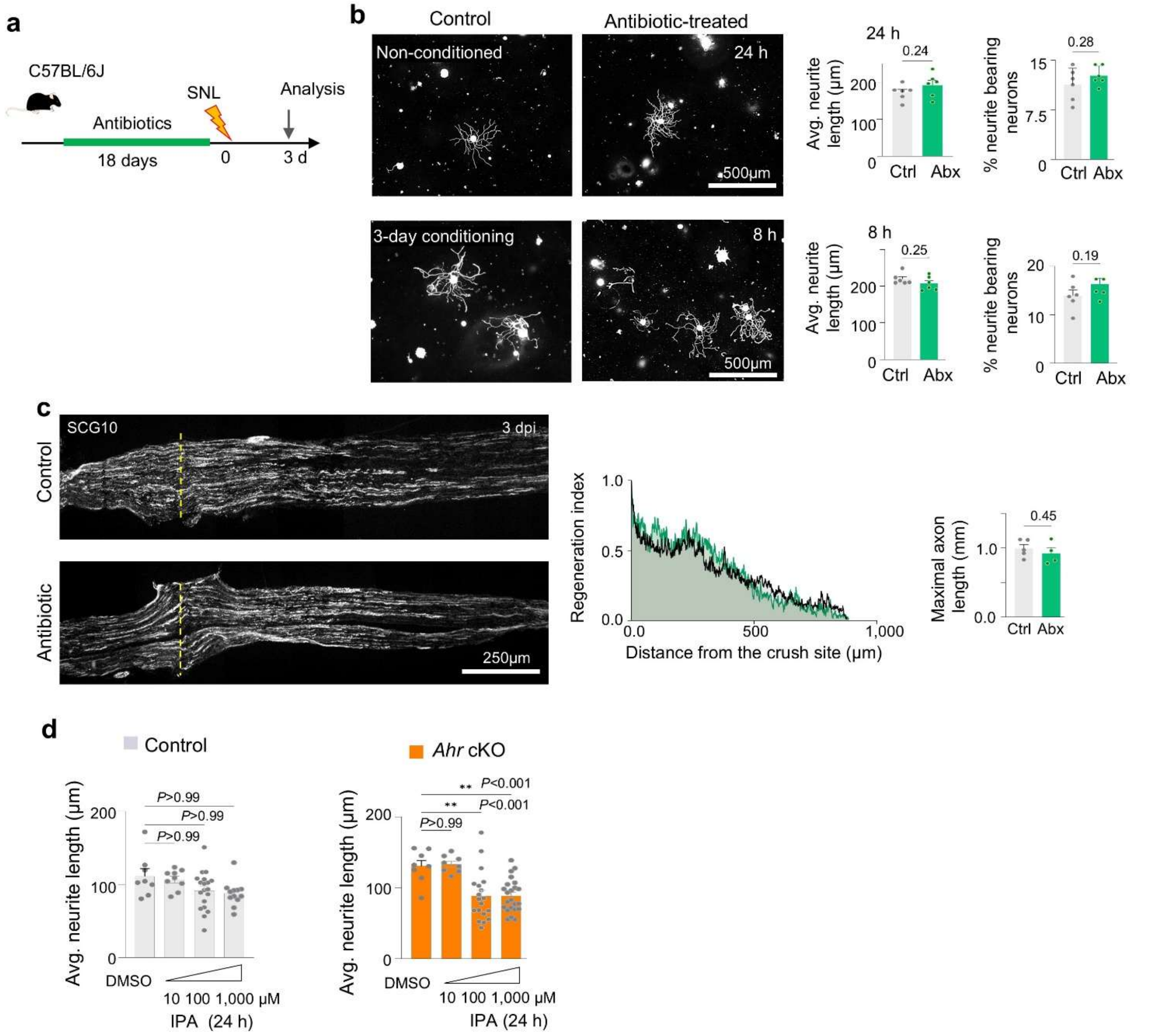


**Figure S7. Antibiotics-induced gut microbiome depletion does not alter axon regeneration after sciatic nerve injury.**

**a.** Experimental paradigm of antibiotics treatment for gut microbiome depletion (18 days pre-treatment before sciatic nerve crush).

**b.** IF images and quantification of neurite length of contralateral and ipsilateral (conditioned) DRG neurons. n = 6 cultures per condition; each data point is mean neurite length from 21-73 neurons. Mean ± s.e.m. Unpaired two-tailed Student's t-test.

**c.** IF images of SCG10$^{+}$ axons at 3 dpi show comparable regeneration in control and antibiotics-treated mice. Dashed lines, lesion center (based on highest SCG10 immunointensity). n = 5 control, n = 4

Abx-treated mice. Regeneration index: two-way ANOVA. Maximal axon length: unpaired two-tailed Student's t-test. Mean ± s.e.m.

**d.** DRG neurons were treated for 24 h with increasing concentrations of AhR ligand indole-3-propionic acid (IPA). Quantifications of mean neurite length of n = 8 to 23 datapoints per condition (each datapoint represents mean from 25 neurons), prepared from 2 mice per genotype. Mean ± s.e.m. One-way ANOVA with Dunnett's correction.

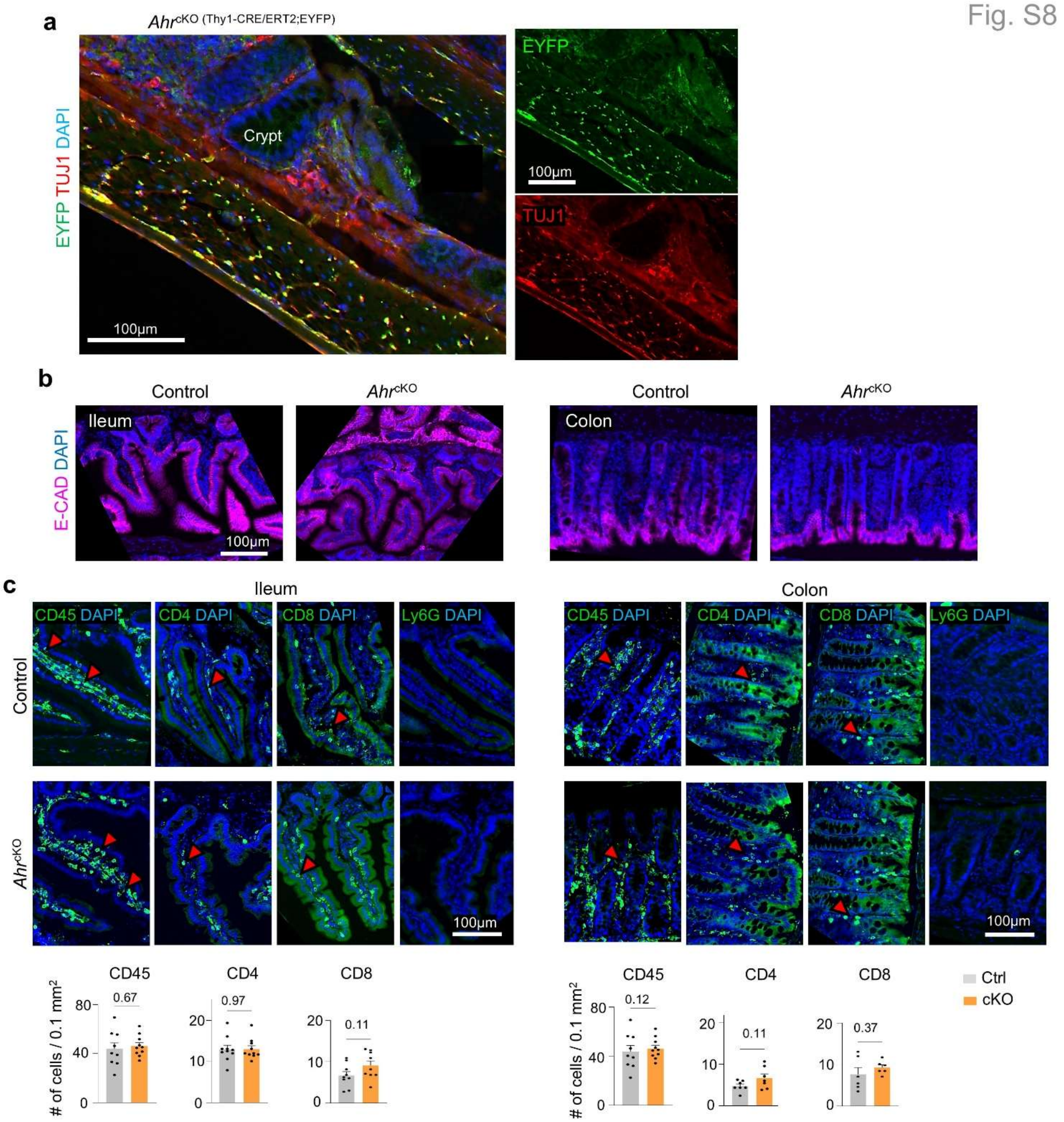


**Figure S8. Neuronal AhR cKO does not alter gut epithelium or immune cell composition.**

**a.** IF images of colonic myenteric plexus in Thy1-CreER/EYFP mice show EYFP expression (by IF with anti-GFP) in $TUJ1^{+}$ enteric neurons.

**b.** IF of distal ileum and colon show intact epithelial architecture in *Ahr*$^{cKO}$ mice compared to wild-type controls. Representative images of n = 3 mice at 1 d post-sham surgery.

**c.** IF and quantification of immune cells in distal ileum and colon from sham-operated cKO and wild-type control mice. Data from 2-3 sections per tissue, n = 3 mice per genotype. Mean ± s.e.m. Unpaired two-tailed Student's t-test.

Fig. S9

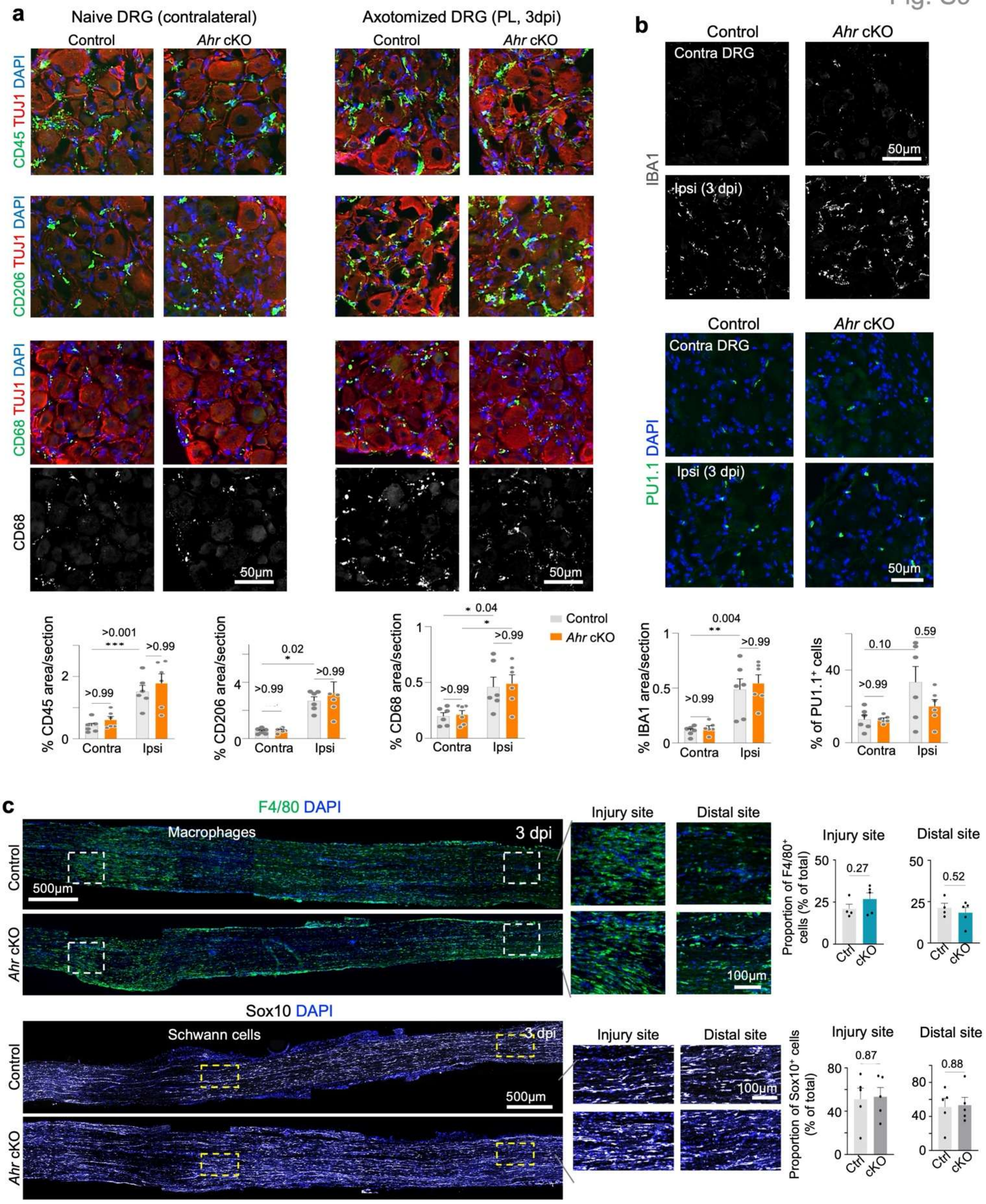


**Figure S9. Neuronal *Ahr* deletion does not affect immune or glial composition in axotomized DRG or injured sciatic nerve.**

**a, b.** IF images and quantifications of immune cell markers in ips- and contralateral DRGs at 3 dpi after

PL. n = 5-6 DRGs from 2 mice per genotype. Mean ± s.e.m. Two-way ANOVA with Bonferroni correction.

**c.** IF staining for F4/80$^+$ macrophages and Sox10$^+$ Schwann cells at the crush site of sciatic nerve at 3 dpi. Quantification of cell proportions at the indicated locations. n = 4 control, n = 5 cKO mice. Mean ± s.e.m. Unpaired two-tailed Student's t-test.

Fig. S10

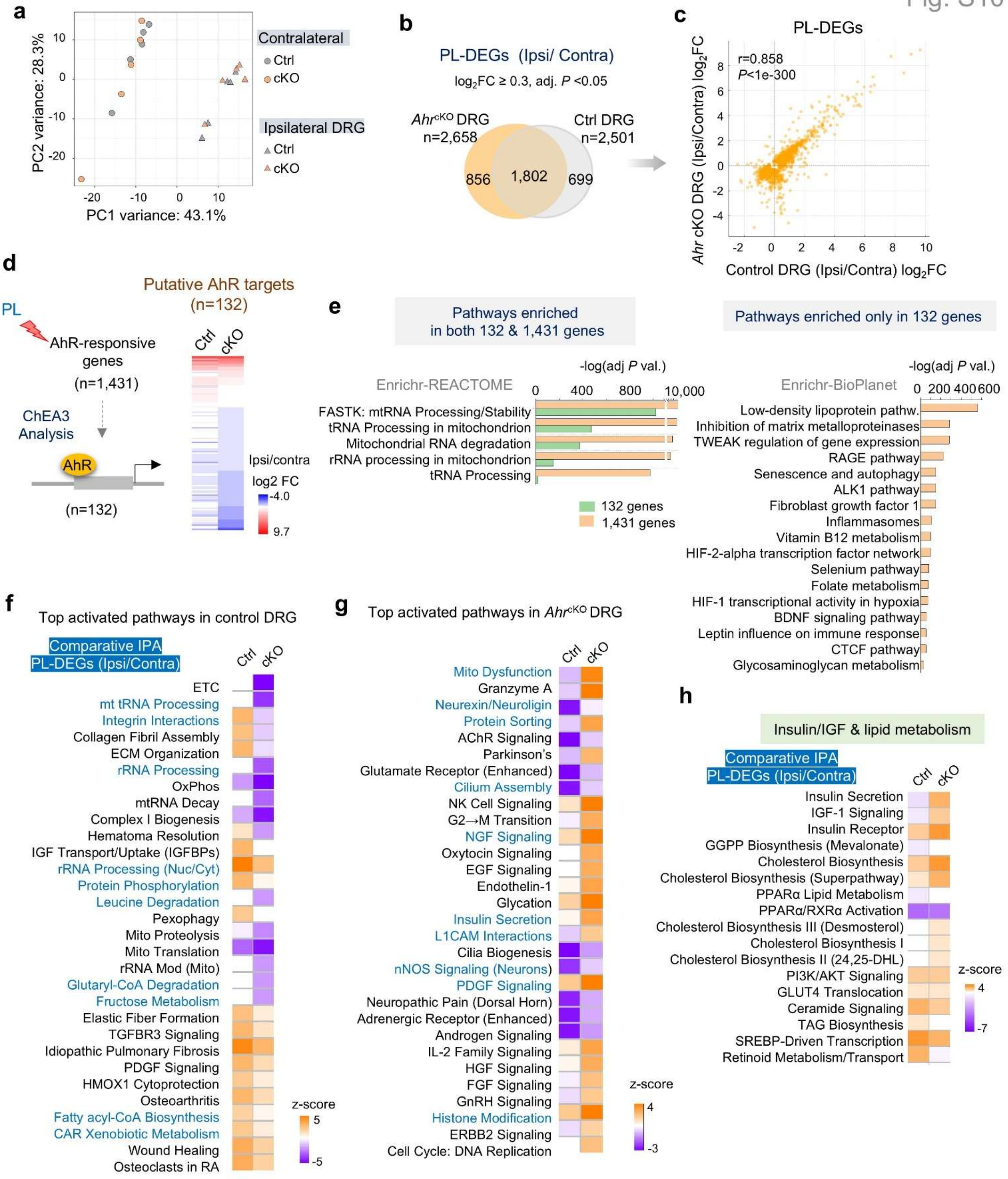


**Fig. S10. Neuronal *Ahr* cKO induces transcriptional shifts in DRG after peripheral lesion.**

**a.** Principal component analysis (PCA) of RNA-seq samples from ipsi- and contralateral sciatic DRGs at 1 dpi (n = 5 per group) shows segregation by injury status, with only modulatory effects of *Ahr* cKO.

**b.** Venn diagram illustrates overlapping and distinct PL-associated genes (ipsi/contra) in cKO vs. controls.

**c.** Scatter plot of $log_2FC$ values for PL-associated genes in control (x axis) versus cKO (y axis).

**d.** Left, ChEA analysis of AhR-responsive genes identified 132 putative AhR target genes. Right, heatmap showing expression ($log_2FC$) of PL-DEGs (ipsi/contra) in control versus cKO.

**e.** Enrichr analysis comparing 1,431 AhR-responsive PL-DEGs to the subset of 132 genes, highlighting shared versus distinctively enriched pathways.

**f, g.** Comparative IPA analyses of ipsi/contra PL-DEGs show the top activated pathways in control versus cKO.

**h.** Comparative IPA of ipsi/contra PL-DEGs reveals differential pathways related to insulin/IGF and lipid metabolism in control versus cKO.

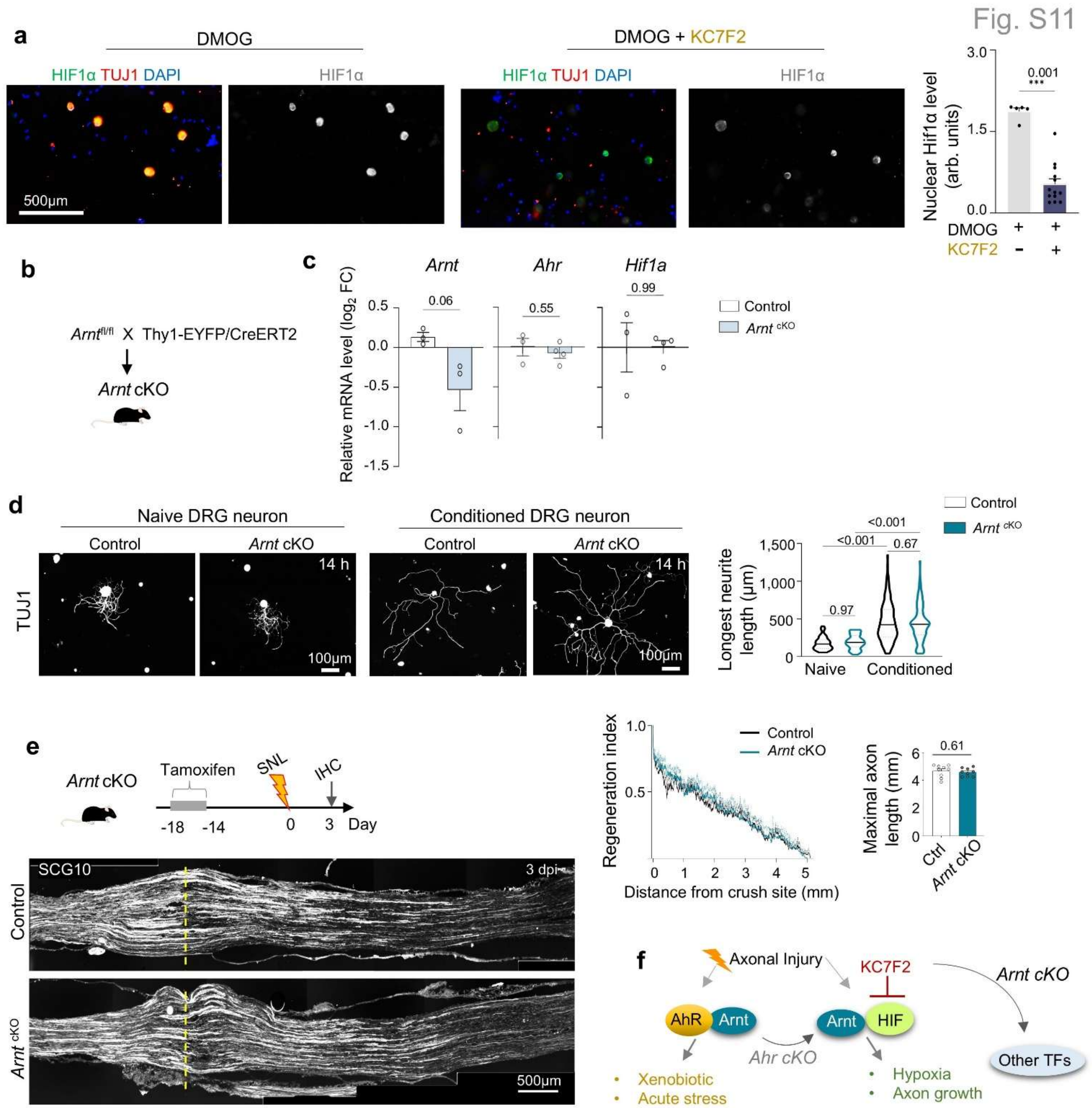


**Figure S11. Neuronal *Arnt* deletion does not impair axon regeneration after sciatic nerve injury.**

**a.** IF of DRG neurons treated with HIF stabilizer DMOG (500 µM) with or without treatment with KC7F2 (80 µM) for 12 h. Quantification of nuclear HIF1α intensity. Mean ± s.e.m. Unpaired two-tailed Student's t-test.

**b.** Generation of *Arnt*$^{fl/fl}$ Thy1-CreERT2/EYFP mice for *Arnt*$^{cKO}$ studies.

**c.** qRT-PCR of *Arnt, Ahr,* and *Hif1a* expression in DRG from *Arnt*$^{cKO}$ and control mice. n = 3-4 per genotype (8 DRGs pooled from 2 mice per sample). Mean ± s.e.m. Unpaired two-tailed Student's t-test.

**d.** IF of TUJ1$^+$ DRG neurons from naive or conditioned DRGs of *Arnt*$^{cKO}$ and control mice at 14 h post-seeding. Violin plots show median and quartiles from n = 51-52 naive, and 122-157 conditioned neurons from 5 mice per genotype. Two-way ANOVA with Bonferroni correction.

**e.** Experimental design of sciatic nerve crush 14 d after tamoxifen injection (100 mg/kg i.p. x 5 d). IF of SCG10$^+$ axons at 3 dpi show comparable regeneration in *Arnt*$^{cKO}$ and control mice. Dashed lines indicate lesion center. n = 9 mice per genotype. Regeneration index: two-way ANOVA. Maximal axon length: unpaired two-tailed Student's t-test.

**f.** Model of AhR-HIF competition for the shared heterodimerization partner ARNT.

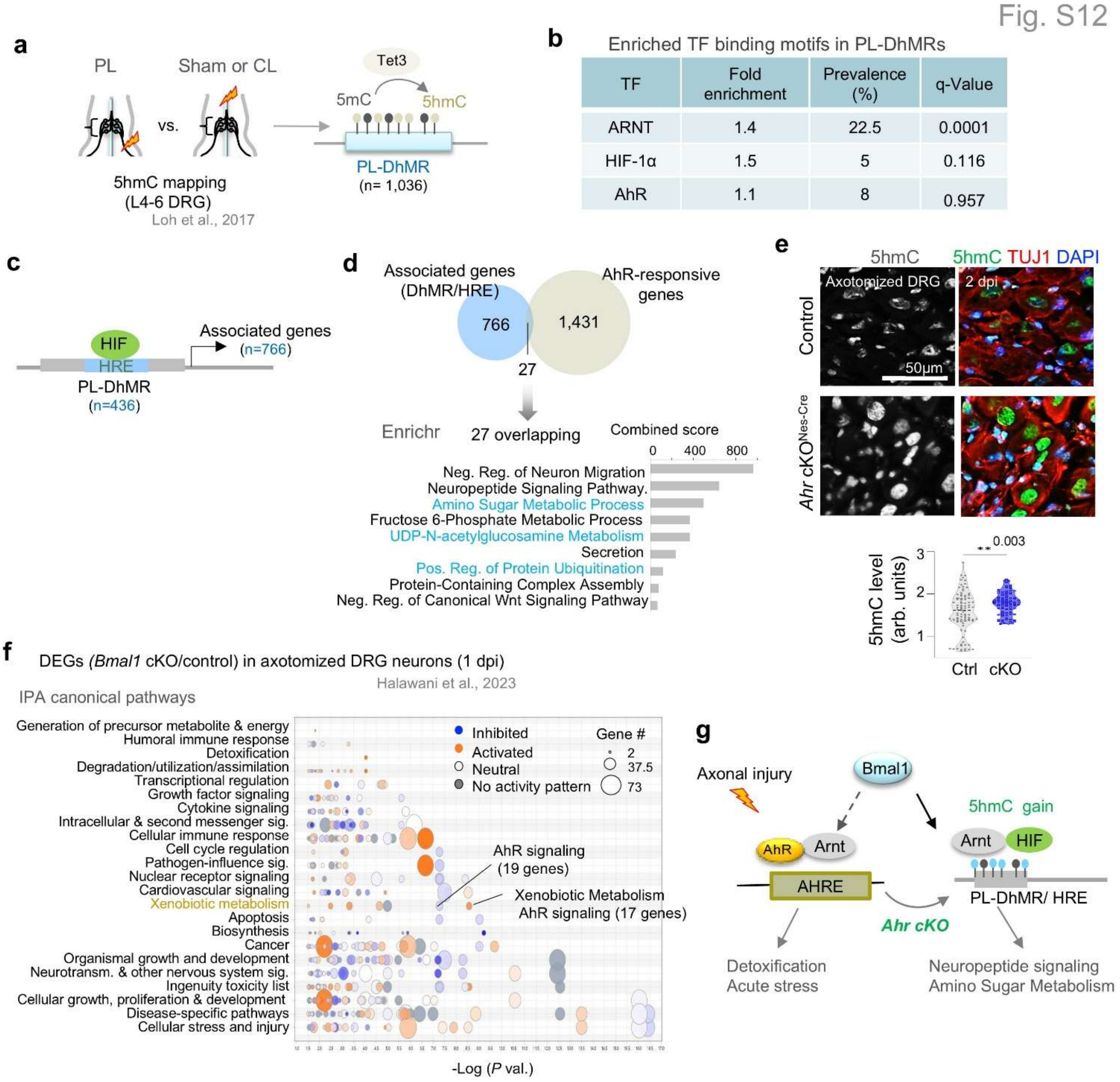


**Figure S12. Interaction of AhR, Bmal1, and DNA hydroxymethylation in response to axotomy.**

**a.** Experimental paradigm for genome-wide mapping of 5hmC after PL in DRG, identifying 1,036 differentially hydroxymethylated regions (DhMRs) compared with sham or central lesion (CL) (Loh et al., 2017). DhMR regulation is mediated by Tet3.

**b.** Summary of HOMER motif analysis of PL-DhMRs (Halawani et al., 2023), showing fold enrichment, prevalence, and FDR (q-value) for ARNT, HIF1α, and AhR motifs.

**c.** Diagram of gene transcription mediated by PL-DhMRs containing HIF response elements (HREs).

**d.** Top, Venn diagram showing overlap of PL-DhMR/HRE-associated genes (n = 766) with AhR-responsive PL-DEGs. Bottom, Enrichr analysis of 27 shared genes highlighting top enriched pathways.

**e.** IF staining and quantification for 5hmC in DRG from *Ahr*$^{cKO}$ (Nestin-Cre) and controls at 2 dpi after PL. n = 9 DRGs from 3 mice per genotype. Median indicated by horizontal line. Mann-Whitney two-tailed test.

**f.** Bubble plot of canonical pathways enriched after *Bmal1* cKO in DRG after axotomy (Halawani et al., 2023), with xenobiotic metabolism and AhR signaling highlighted.

**g.** Working model. Arnt-HIF1α complex may act at 5hmC-modified loci in DRG neurons to regulate pro-regenerative programs after axotomy. *Ahr* deletion skews ARNT activity toward the HIF1α regulon, partly gated by Bmal1-dependent 5hmC modifications.

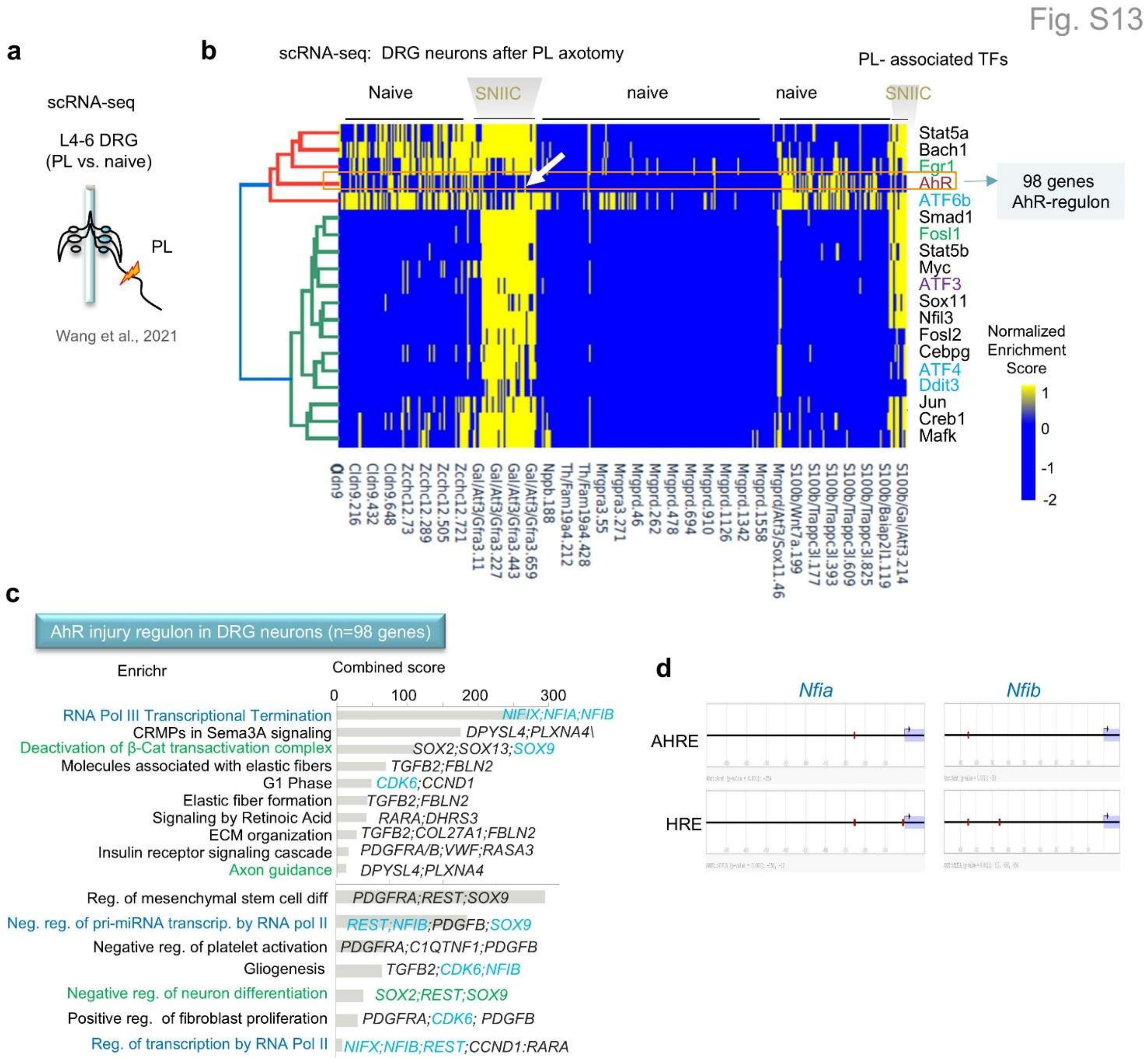


**Figure S13. scRNA-seq of DRG reveals neuronal AhR injury regulon associated with RNA regulation and integrated stress response.**

**a.** Schematic of scRNA-seq of axotomized sciatic DRG after PL compared to naive DRG (Wang et al., 2021).

**b.** Heatmap of normalized enrichment scores for PL-associated transcription factors in DRG neurons after axotomy (Wang et al., 2021). Sciatic nerve injury-induced neuronal clusters (SNIIC), defined by ATF3 expression, showed AhR in OFF state (white arrow), in contrast to ON state for majority of TFs, including ATF3 and regulators of Integrated stress response (ISR; denoted in blue).

**c.** Enrichr pathway analysis of a predicted AhR regulon of 98 genes in axotomized DRG neurons, showing enrichment in RNA processing and ISR-related pathways in blue.

**d.** Eukaryotic Promoter Database analyses show AHRE and HRE motifs in promoters of *Nfia* and *Nfib* genes.

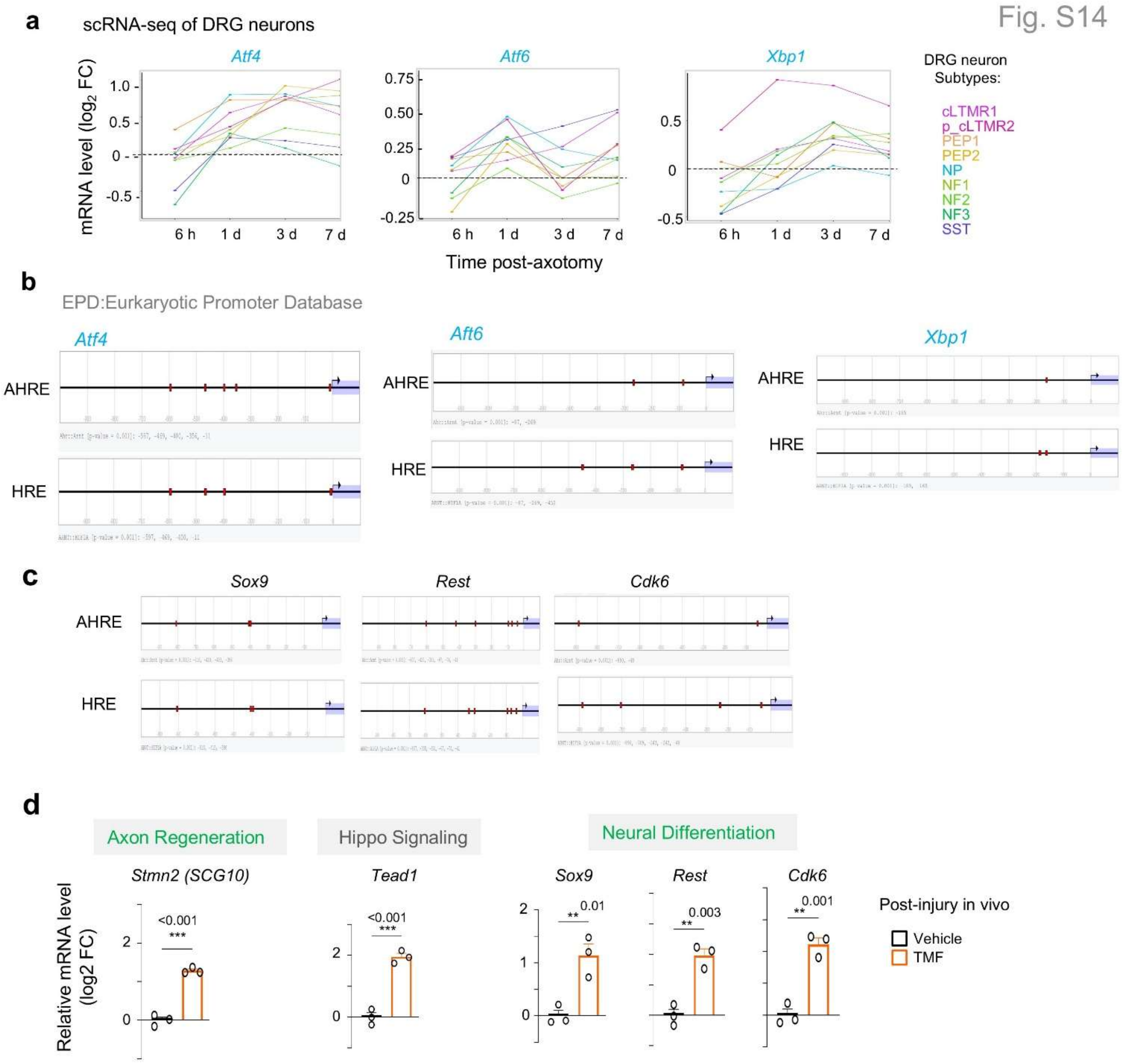


**Figure S14. Stress response transcription factors induced by axotomy.**

**a.** scRNA-seq survey of conditioned DRG (in dataset Renthal et al., 2019) shows induction of *Atf4, Atf6*, and *Xbp1* in the majority of DRG neurons after PL.

**b, c.** Eukaryotic Promoter Database analyses show AHRE or HRE motifs in promoters of the indicated genes.

**d.** qRT-PCR expression analysis of selected genes linked to neural differentiation and axonogenesis. *Hprt1* expression used for normalization. n = 3 independent samples (each pooled from L4-L6 DRGs from 3 mice per group). Mean ± s.e.m. One-way ANOVA with Dunnett's correction.
